# Experience-dependent reorganization of inhibitory neuron synaptic connectivity

**DOI:** 10.1101/2025.01.16.633450

**Authors:** Andrew J.P. Fink, Samuel P. Muscinelli, Shuqi Wang, Marcus I. Hogan, Daniel F. English, Richard Axel, Ashok Litwin-Kumar, Carl E. Schoonover

**Affiliations:** Department of Neurobiology, Northwestern University Evanston, IL; Mortimer B. Zuckerman Mind Brain Behavior Institute Department of Neuroscience Columbia University New York, NY; École Polytechnique Fédérale de Lausanne Lausanne, Switzerland; Neuroscience Graduate Program, University of California Berkeley Berkeley, CA; School of Neuroscience Virginia Tech Blacksburg, VA; Howard Hughes Medical Institute; Allen Institute for Neural Dynamics Seattle, WA

## Abstract

Organisms continually tune their perceptual systems to the features they encounter in their environment^1–3^. We have studied how ongoing experience reorganizes the synaptic connectivity of neurons in the olfactory (piriform) cortex of the mouse. We developed an approach to measure synaptic connectivity *in vivo*, training a deep convolutional network to reliably identify monosynaptic connections from the spike-time cross-correlograms of 4.4 million single-unit pairs. This revealed that excitatory piriform neurons with similar odor tuning are more likely to be connected. We asked whether experience enhances this like-to-like connectivity but found that it was unaffected by odor exposure. Experience did, however, alter the logic of interneuron connectivity. Following repeated encounters with a set of odorants, inhibitory neurons that responded differentially to these stimuli exhibited a high degree of both incoming and outgoing synaptic connections within the cortical network. This reorganization depended only on the odor tuning of the inhibitory interneuron and not on the tuning of its pre- or postsynaptic partners. A computational model of this reorganized connectivity predicts that it increases the dimensionality of the entire network’s responses to familiar stimuli, thereby enhancing their discriminability. We confirmed that this network-level property is present in physiological measurements, which showed increased dimensionality and separability of the evoked responses to familiar versus novel odorants. Thus, a simple, non-Hebbian reorganization of interneuron connectivity may selectively enhance an organism’s discrimination of the features of its environment.

## Main text

Olfactory perception is initiated by the binding of odorants to receptors on primary sensory neurons in the olfactory epithelium^4^, which in turn send axonal projections to glomeruli in the olfactory bulb^5^. Olfactory information is then conveyed to the olfactory (piriform) cortex by the bulb’s mitral and tufted cells^6^. In the anterior piriform synaptic connections between pyramidal neurons are highly distributed: the vast majority of inputs to a neuron are from distal partners^7–9^, resulting in a network where connection probability is low (∼0.1%) even locally^6,8^. This sparse connectivity limits the probability that a pair of piriform neurons is driven by a common presynaptic partner, a confound that has historically hindered inference of monosynaptic connections from neuronal spiking. Thus, the piriform presents an opportunity to observe both the spiking activity of a population of neurons and their synaptic connectivity in an awake, behaving mouse. We have studied the relationship between piriform activity and connectivity and how this relationship is modified by experience.

### Reliable inference of monosynaptic connections

We performed recordings in the anterior piriform cortex of awake, head-fixed mice using a 4-shank Neuropixels silicon probe^11^. The cell-dense layer in the anterior-most portion of the cortex folds over itself (**Extended Data Fig. 1a**), spanning a substantial fraction of our probe’s electrode sites. This anatomical feature permitted us to obtain 783 ± 128 single units per recording (mean ± s.d.; range, 642 - 1,007; N = 7 mice), the vast majority of which were located in this single brain region. Given the sparse connectivity of the piriform network, such a high yield was critical to identify a sufficient number of connected pairs.

We inferred monosynaptic connectivity from spike-time cross correlograms^12^ computed using the spontaneous activity of 4,401,362 single-unit pairs (**Fig. 1a,b**). We developed a deep convolutional network (Dyad, **Extended Data Fig. 1c**) to identify those pairs whose correlograms had a peak consistent with an excitatory monosynaptic interaction: a sharp rise at short latency followed by a slower decay^13–20^ (**Fig. 1a,c, Extended Data Fig. 1b**). Given the inherent difficulty in distinguishing synaptically connected pairs from pairs receiving common drive from a third neuron^12,13,18,21^, we validated Dyad using a previously published ground truth dataset^22^ with positively identified monosynaptic excitatory connections *in vivo* (see **Methods**). We computed the proportion of inferred connections that corresponded to true synapses (precision) and the proportion of inferred connections over the total number of true synapses (recall). In this dataset Dyad identified excitatory synapses without error up to a recall of 47% (**Fig. 1d**), perfectly recovering approximately half of the synapses. This indicates that connections inferred by Dyad very likely reflect true synaptic contacts. We note, however, that this ground truth dataset contains only thirty connected pairs.

**Figure 1.**
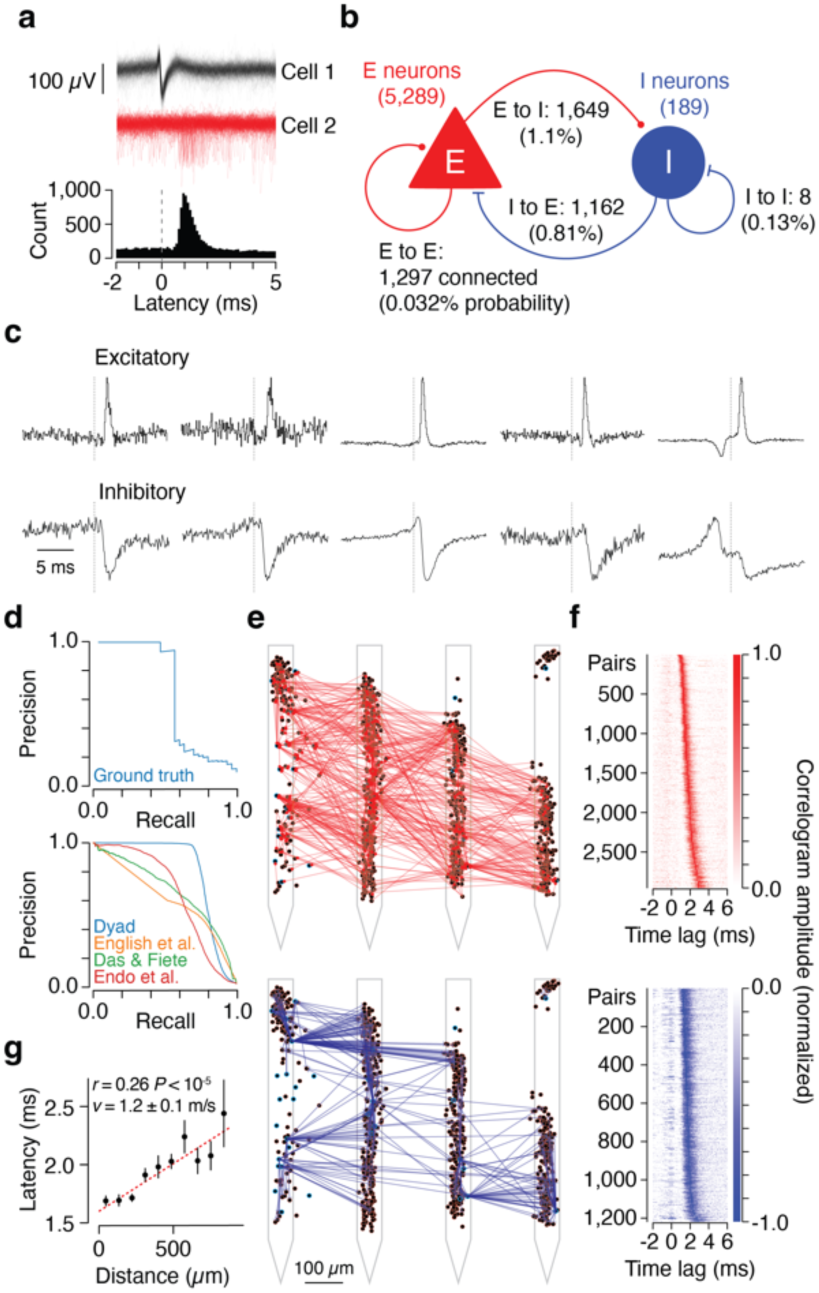
Reliable inference of monosynaptic connections in the piriform cortex. **a**, Spike waveforms of a pair of cells (black and red traces, top), aligned to the spike times of cell 1. The increased spiking probability of cell 2 after a cell 1 spike is captured in the spike-time cross-correlogram (CCG) of the pair (bottom), which measures the probability that cell 2 fires an action potential at a range of time lags relative to cell 1. For a synaptically connected pair of neurons, a presynaptic spike produces an asymmetric peak on the causal side of the correlogram, whose shape reflects the underlying excitatory postsynaptic potential: a sharp rise at short latency followed by a slower decay to baseline^13–20^. Precise measurement of this shape requires several hours of continuously recorded spike trains (6.1 ± 0.9 hours, mean ± s.d., min = 5.3 hours, max = 7.1 hours). **b**, Summary of the number of recorded single units and inferred connections (and corresponding connection probabilities). E to E: 1,297 connected out of 4,107,263 total pairs from 7 recordings; E to I: 1,649 connected out of 144,056 pairs; I to E: 1,162 connected out of 144,056 pairs; I to I: 8 connected out of 5,798 pairs. **c**, Examples CCGs for excitatory (top) and inhibitory (bottom) monosynaptic connections identified by Dyad. The two rightmost examples correspond to reciprocally connected E-I pairs. **d**, Dyad validation. Top: Precision-recall curve on an *in vivo* ground-truth dataset with positively identified excitatory synaptic connections^22^. Bottom: Precision-recall curve on simulated ground-truth data (see **Methods**) for Dyad (blue) and previously published connectivity inference methods: English^22^ (orange, peak-detection algorithm); Das^21^ (green, GLM-based inference); Endo^23^ (red, deep learning approach). These results are for a simulated network with 2.5% connection probability. **e**, Estimated single unit locations (black circles) and inferred connectivity excitatory (red lines, top) or inhibitory (blue lines, bottom) for one example dataset. The contour of the circle indicates whether the single unit was classified as an E neuron (red) or an I neuron (blue). **f**, Heatmap of normalized CCGs for all pairs connected by an excitatory (top) or inhibitory (bottom) synapse identified by Dyad. CCGs were normalized for visualization purposes. **g**, Peak latency as a function of distance between the pair. Black markers: mean latency ± 95% confidence intervals at 10 equally-spaced bins of distance; red dashed line: linear fit to all the connected pairs (i.e. prior to binning). *r,* P: Pearson’s correlation coefficient and corresponding P-value. (N = 2,946 pairs) The axon conduction velocity *v* was estimated from the slope of the red dashed line.

We therefore next evaluated the ability of Dyad to identify excitatory monosynaptic connections from simulated spike times produced by a network consisting of recurrently connected leaky integrate and fire neurons (**Extended Data Fig. 1d)**. We found that Dyad’s precision exceeds 0.99 up to a recall of 0.63 for a sparsely connected network (**Fig. 1d**). Thus Dyad recovered more than two thirds of the network’s synapses with near-perfect accuracy and outperformed state-of-the-art approaches^21–23^ for inferring synaptic connectivity from spike trains (**Fig. 1d**). Both precision and recall increased with the number of spikes in our simulations, emphasizing the importance of long recordings (**Extended Data Fig. 1f**). Consistent with prior theoretical work^24,25^, we observed that precision also improved with recurrent network sparseness (**Extended Data Fig. 1e, g**). As connection probability falls, so does the prevalence of common inputs to a pair of neurons as well as polysynaptic chains, both of which introduce correlations in spiking that confound estimation of monosynaptic connectivity^21,24–26^. Thus, the piriform is likely to support reliable synapse inference owing to its exceptionally sparse recurrent connectivity.

We employed Dyad to detect putative excitatory monosynaptic connections in our piriform recordings and obtained 2,946 connected pairs, or 0.067% of all pairs (**Fig. 1b,e,f**). This inferred connection probability is lower than anatomical estimates (∼0.1%), in agreement with our simulations and analysis of ground-truth data in which Dyad is conservative and recovered only a fraction of all connections at high precision. We devised three independent analyses to estimate the rate of error in our piriform recordings (see **Extended Data Fig. 1h,j,k**). In the first analysis, we counted how often Dyad spuriously inferred an excitatory connection arising from a likely inhibitory neuron (see **Methods**). We found that Dyad identified only 2 such pairs out of 148,689 total possible interactions (**Extended Data Fig. 1h**), indicating a false positive rate of approximately 1.3 x 10^-5^ and an estimated precision ∼98% (see **Methods**). In the second analysis we determined the likelihood of finding correlograms with a sharp, asymmetric peak whose onset preceded rather than followed the zero time lag. Such a peak would necessarily be caused by common input rather than the spiking of the presynaptic neuron. We found that these spurious peaks occurred with a false-positive rate of 8.1 x 10^-6^ (**Extended Data Fig. 1j**, see **Methods**), providing a second, independent estimate of our precision >98%. Finally, we asked whether the connections we inferred reflected poly-rather than mono-synaptic input. We studied all sets of three single units in our data that Dyad identified as connected in a disynaptic chain. We then asked how often Dyad inferred a synaptic connection between the first and last single unit in these triplets. A detection rate greater than that expected by chance would imply spurious inferred connections due to disynaptic chains. However, Dyad identified such connections at a rate indistinguishable from chance when controlling for the connectivity statistics of the network (in-degree and out-degree) (**Extended Data Fig. 1k**). Therefore, the connectivity we infer overwhelmingly reflects mono-rather than polysynaptic interactions. We note that weak connections, or connections onto postsynaptic neurons that have low input resistance or low firing rates are those most likely to evade our detection (**Extended Data Fig. 1i**). Nonetheless, we observe a broad distribution of efficacies indicating that our method can recover connections that span a wide range of strengths (**Extended Data Fig. 1l)**.

We also developed a similar approach for inferring inhibitory interactions. We trained a second deep neural network to find pairs of single units whose correlograms exhibit a sharp, downward peak at short latency, consistent with the presence of an inhibitory synapse^12,13,18,19,22,27^ (**Fig. 1c,e,f**). We validated this approach using strategies similar to those applied above, although note that there is no comparable ground truth data set for inhibitory connections. We found that this second neural network identified inhibitory connections with a precision > 99% when considering only fast-spiking single units (**Extended Data Fig. 1h,j**, see **Methods**). However, validation of this method on recurrent network simulations showed a lower recall than our inference of excitatory connectivity (precision > 99% up to recall of ∼ 55%, **Extended Data Fig. 1g,** see **Methods**).

Finally, we asked whether Dyad successfully recovers key known properties of our circuit. We segregated the population into putative inhibitory neurons (fast-spiking single units, “I neurons”), and putative excitatory neurons (regular-spiking single units, “E neurons”) (**Extended Data Fig. 1h**, see **Methods**). First, we considered the sign of inferred connections and confirmed that I neurons tend to inhibit, and E neurons tend to excite their postsynaptic partners^27^ (**Extended Data Fig. 1h**). Second, we assessed whether the latency to peak in excitatory correlograms (1.8 ± 0.5 ms, mean ± s.d.) scales with the distance between the pair (**Fig. 1g, Extended Data Fig. 1m**). The slope of this relationship yielded an estimate of axon conduction velocity of 1.2 ± 0.1 m/s (mean ± s.e.m., Pearson’s *r* = 0.26, R^2^ = 0.068, N = 2,954 pairs, **Fig. 1g**), in agreement with direct measurements of conduction velocity in this cortex^28^. Third, we found that I neurons that receive many connections also tend to form many outputs (Pearson’s *r* = 0.62, P < 10^-10^, N = 189 neurons, **Extended Data Fig. 1n**). We confirmed this correlation of inputs and outputs by quantifying the connectivity of basket cells in a volume electron microscopy reconstruction^29^ (**Extended Data Fig. 1o**). From the morphology and connectivity of these basket cells^30,31^, they most likely correspond to parvalbumin-positive fast-spiking inhibitory interneurons. We found a strong correlation between the number of excitatory inputs onto a basket cell and the number of inhibitory outputs from that cell onto excitatory neurons (Pearson’s *r* = 0.79, P = 4.49 x 10^-13^, N = 57 neurons from one mouse). Thus, our method for synapse inference recovers basic biophysical and anatomical properties. We conclude that Dyad detects excitatory and inhibitory monosynaptic connections with high confidence but recovers only a subset of synapses between the recorded neurons.

### Like-to-like connectivity in piriform

We next employed Dyad to examine the relationship between odor tuning and synaptic connectivity. We recorded piriform activity in response to eight chemically and perceptually distinct odorants of neutral valence (**Fig. 2a, Extended Data Fig. 2**). We presented each odorant 25 times each to awake, head-fixed mice that had not received any prior experience with the stimuli. Consistent with prior reports, we observed odor-selective evoked responses, including in I neurons^32–37^ (**Extended Data Fig. 2p-x**). We inferred connectivity from stretches of spontaneous activity and measured odor responses during separate portions of the recording. This revealed a like-to-like connectivity motif: E-E neuron pairs that responded similarly to the panel of odorant stimuli exhibited enriched connectivity (**Fig. 2b,** P = 2.8 x 10^-10^, null models described in figure legend). The high precision with which Dyad recovers monosynaptic connectivity implies that this like-to-like motif is unlikely to reflect common input from the olfactory bulb. This organization resembles reports of superficial pyramidal neuron connectivity in the visual cortex of the mouse^38–40^. In the piriform this basic pattern additionally holds for E-to-I and I-to-E pairs (**Extended Data Fig. 3b,c,** P < 0.001 for all panels). This stands in contrast to the visual cortex, where like-to-like connectivity between pyramidal and parvalbumin-positive inhibitory interneurons has been reported for connection strength but not connection probability^41^. This difference might be explained in part by our method’s bias towards strong synapses.

**Figure 2.**
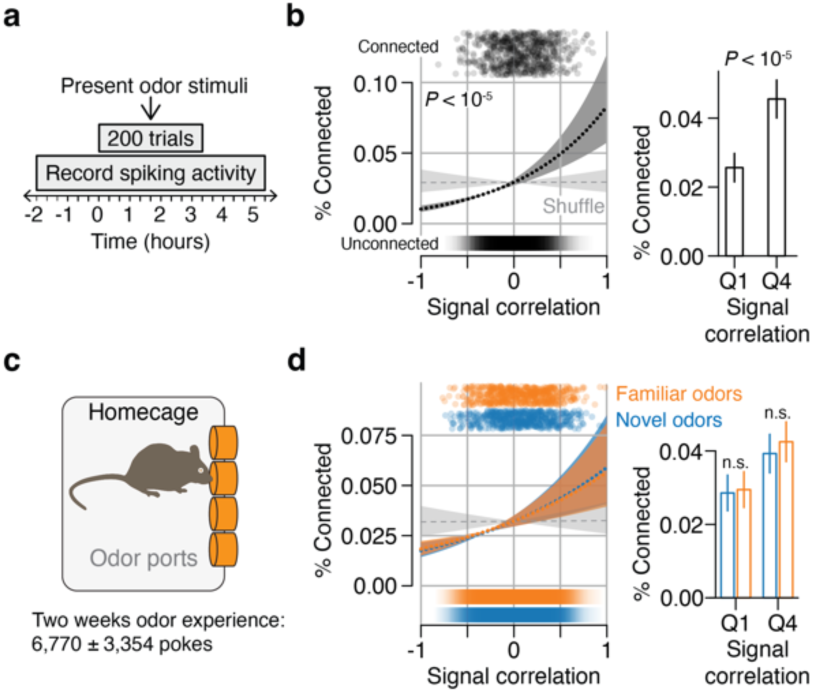
Like-to-like connectivity in the piriform is not enhanced by experience. **a**, Recording session timeline. The first odor stimulus (t = 0) is presented ∼2 hours after the start of the recording. Eight odorant stimuli were presented in 25 blocks, in pseudorandom order in each block. Each odor was delivered for 4 seconds, with 1 minute inter-stimulus interval. After the last odor stimulus, spontaneous activity was recorded for additional ∼2 hours. **b**, Left: probability of E-to-E connections as a function of signal correlation (see **Methods**). Black markers at the top show the signal correlation of connected pairs, jittered among the y-axis for visualization purposes. The heatmap at the bottom shows the density of unconnected pairs. Dotted line: logistic regression fit; shaded area: 95% confidence interval. Gray dashed line and shaded area: logistic regression on a null model obtained by shuffling odor stimulus identities independently for each E neuron. This null model was used to compute both P-values in this panel. Right: mean connection probability and 95% confidence intervals (Clopper-Pearson) of pairs in the lower and top quartiles of signal correlation (148 and 263 connected pairs in the lower and top quartile respectively, out of 577,376 pairs). **c**, Odor familiarization paradigm^45^. Four odor ports (orange) were inserted into the wall of a modified homecage. Animals underwent this familiarization protocol for approximately two weeks, then recorded from as in **a**. The panel of odorant stimuli consisted of the four familiar odorants that were administered in the home cage along with four novel stimuli. **d**, Same as **b**, but for experienced animals and where signal correlations were computed either across novel (blue) or familiar (orange) stimuli. Significance was tested with respect to a null model in which we constructed pseudo-novel and pseudo-familiar odor sets, each containing two random odors from the novel set and two from the familiar set.

It has been proposed that like-to-like connectivity is assembled by an experience-dependent, Hebbian process^42–44^. However, to our knowledge the odors we employed were novel to the animals. We therefore asked whether this like-to-like connectivity may have emerged from spatial organization in the piriform network. Indeed, connection probability was higher for nearby neurons, falling off steeply over the first 200 μm (**Extended Data Fig. 3d–f**). Above 200 μm, connection probability was approximately three-fold lower, exhibiting the slow spatial decay consistent with previous reports at these distances^8,9^. We also observed a weak dependence of response similarity on distance (**Extended Data Fig. 3g,h**). We reasoned that like-to-like connectivity might emerge without experience if nearby neurons are both more likely to be connected to each other and more similar in their tuning. However, a model in which connectivity was shuffled while preserving these distance dependencies failed to reproduce the data (**Extended Data Fig. 3i–k**). We conclude that the spatial structure we detect is not sufficient to explain like-to-like connectivity in the piriform.

### Experience reorganizes piriform connectivity

We therefore considered the hypothesis that like-to-like connectivity arises from experience-dependent plasticity^42,43^ and asked whether extensive exposure to odorants can further increase the probability of connections between similarly tuned neurons. This would be consistent with a Hebbian mechanism enhancing synaptic coupling between correlated neurons that are repeatedly co-activated by sensory stimuli^44^. Mice were afforded extensive experience with four neutral odorants over a period of two weeks: four odor ports were placed in the walls of the animals’ home cages such that they could sample the stimuli at will^45^ (**Fig. 2c, Extended Data Fig. 4a**). The animals had *ad libitum* access to food and water, and the stimuli were not explicitly reinforced. On average, the mice volitionally sampled each odor more than a thousand times over this two-week period (6,770 ± 3,354 samples, mean ± s.d.; range, 1,209 - 9,989 samples, **Extended Data Fig. 4a, center**). We then recorded from these mice (“experienced animals”) and presented eight odors: the four familiar odors they had sampled over the previous two weeks and four novel odors. We performed the same analysis on data from the experienced animals as we had previously for the naïve cohort. Once again, we found that, for novel stimuli, connection probability scaled with response similarity. However, we did not detect any enhancement of this relationship for the familiar odors, as would have been expected from a Hebbian process (**Fig. 2d, Extended Data Fig. 4b–d**). We note that our conclusions are limited by the sensitivity of our assay: we do not detect all of the synaptic connections between the recorded neurons; and experience may alter synaptic weights in a way that we cannot observe since Dyad reports binary connections not strengths. It is also possible that the formation of Hebbian ensembles in the piriform requires odors to be paired with reinforcement^46^. However, our results demonstrate that regular experience with odorants does not measurably accentuate like-to-like connectivity in the adult piriform.

We next examined whether experience alters the relationship between connectivity and response properties other than co-tuning. Specifically, we considered the properties of individual neurons rather than the relationship between pairs of neurons. We asked whether connection counts between E-to-E and E-to-I pairs covary with differences in evoked response amplitude, trial-to-trial variability, and odor tuning for novel versus familiar sets of stimuli. After correcting for multiple comparisons, we found only two significant effects. First, we found a weak (Pearson’s *r* = 0.06) relationship between the coefficient of variation of the odor response of E neurons across trials and their number of outgoing connections (**Extended Data Fig. 5f**). We also found a strong effect of experience on the relationship between the tuning of an I neuron and its connectivity, which we explore further (**Fig. 3, Extended Data Fig. 5**).

**Figure 3.**
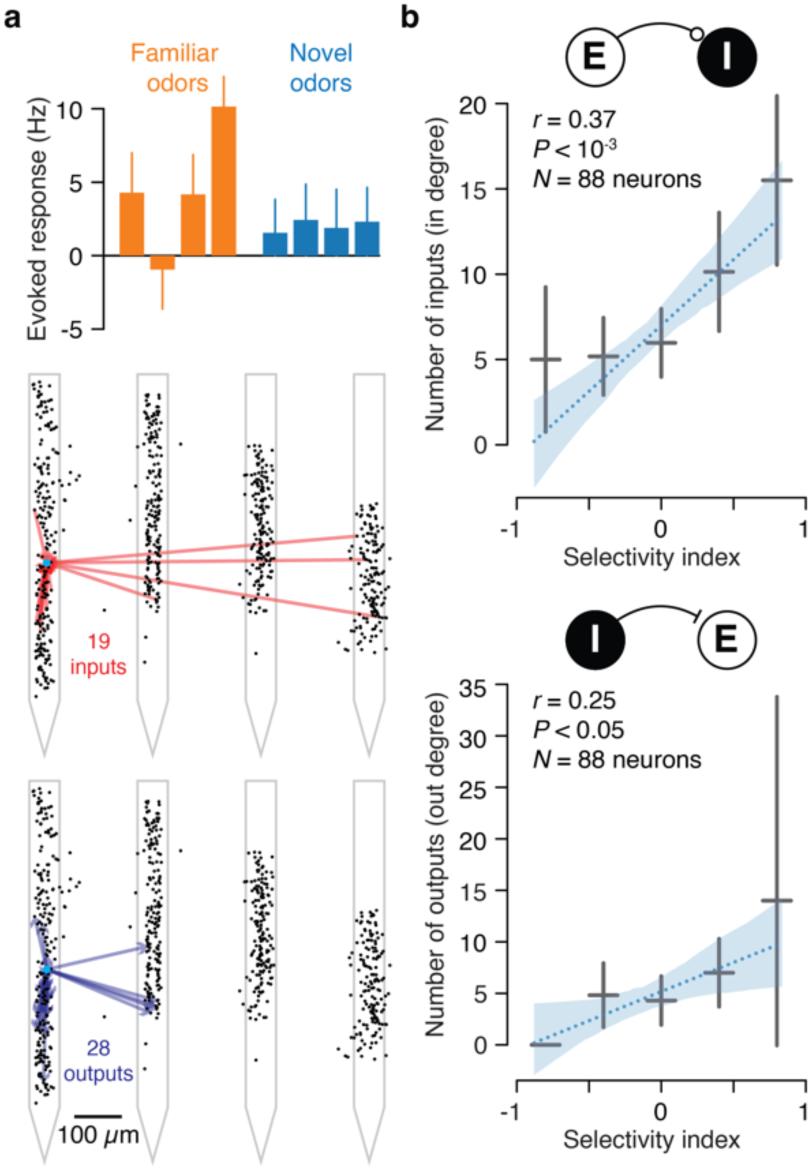
Effect of experience on the connectivity of inhibitory interneurons. **a,** Example I neuron in an experienced animal. Top: evoked response amplitudes for the 4 familiar and 4 novel odors (trial average ± s.e.m.); bottom: connectivity (black circles: estimated unit locations; inferred excitatory inputs: red arrows; inferred inhibitory outputs: blue arrows; blue circles: estimated position of example I neuron). **b,** In degree (top) and out degree (bottom) of I neurons in experienced animals as a function of the I neurons’ index (SI). Black markers: the means and 95% confidence intervals in equally-spaced bins of SI. Blue dotted line: linear fit to the individual data points (i.e. prior to binning); blue shaded region: 95% confidence interval on this fit. P-values were obtained with respect to a null model in which S.I. and in/out degrees were independently shuffled 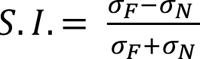. where σ_*N*_, σ_*F*_ are the standard deviation of the trial-averaged odor response across novel or familiar odors, respectively (see **Methods**).

In experienced animals, I neurons that responded more differentially across familiar odors also received more excitatory inputs and produced more inhibitory outputs (**Fig. 3a, Extended Data Fig. 5**): we found a strong correlation between an I neuron’s selectivity across familiar stimuli and its number of inputs (in-degree, **Fig 3b, top**) and outputs (out-degree, **Fig 3b, bottom**). This pattern held when considering the sum of in- and out-degrees for each I neuron (**Extended Data Fig. 5c, right**), consistent with our observation that the number of inputs (in-degree) and outputs (out-degree) are correlated in I neurons (**Extended Data Fig. 1n**). Moreover, high-degree I neurons (normalized degree > 0.015) were more differentially modulated by familiar odors than low-degree I neurons (normalized degree < 0.015) (**Extended Data Fig. 5d**). Thus, in experienced mice, the I neurons most differentially responsive across odors in the animal’s recent history also tend to be densely interconnected in the piriform network.

We asked whether this organization is experience dependent by performing the same analyses in our cohort of naïve mice, who had no prior experience with the “familiar” and “novel” odor sets. Each of the dependencies we observed in experienced mice differed significantly from those found in naïve animals, where in all cases we found no correlation between degree and odor selectivity (**Extended Data Fig. 5c, bottom**), including when we considered all possible ways to divide the 8 novel odors in two sets (**Extended Data Fig. 5e**). Moreover, in naïve animals the selectivity of both high- and low-degree I neurons did not differ significantly (**Extended Data Fig. 5d**). Therefore, the relationship we observe between the connectivity and the tuning of I neurons depends on experience.

Notably, we detected no difference between naïve and experienced animals in the distributions of both degree and of odor selectivity across I neurons (**Extended Data Fig. 6**). Therefore, any gains in connectivity or selectivity in a subset of I neurons must be balanced by a proportional loss in other cells. These results suggest that in naïve animals, highly connected I neurons may be highly selective across odors in the animals’ home environment, rather than the novel stimuli we administered during our recordings.

These population phenomena are robust (**Extended Data Fig. 7**). They were observed in each individual mouse (**Extended Data Fig. 7a**) and were only weakly sensitive to the choice of quantification window (**Extended Data Fig. 7b**) and synapse inference inclusion threshold (**Extended Data Fig. 7e**). We considered the possible confound that experience increases the excitability of selective I neurons, enhancing the detection of afferent synapses and consequently producing a spurious dependence of in-degree on selectivity. However, the firing rates of I neurons did not covary with selectivity (**Extended Data Fig. 7c**). We found only negligible effects of experience on the basic response properties of the piriform, including evoked firing rate, lifetime sparseness, population sparseness, and the sharpness of tuning (**Extended Data Fig. 8**). We also did not observe an effect of experience on Layer 1 feedforward interneurons or on likely Somatostatin-positive feedback interneurons but note that these were rare in our recordings (8 and 15 single units, respectively, **Extended Data Fig. 7d**). Finally, we determined that these findings are unlikely to reflect spike sorting contamination for nearby neurons, as all of the results (**Fig 2** and **Fig. 3**) held when considering only connections between neurons recorded on different probe shanks (**Extended Data Fig. 9**).

Interestingly, for E/I pairs, neither the pre- nor the postsynaptic E neurons’ selectivity covaried with connection probability (**Extended Data Fig. 7f,g)**. Thus, the experience-dependent reorganization of the network is a sole function of the I neuron’s selectivity and does not appear to obey Hebbian principles. This reordering of the relationship between connectivity and tuning may reflect an experience-dependent plasticity rule under which the selectivity of an I neuron to the odors in the environment determines the number of its inputs and outputs. Alternatively, experience may not alter the wiring of I neurons; rather the most densely connected ones may become more differentially responsive to the experienced odors.

### Effect of inhibitory wiring on network function

Finally, we examined how this dependence between I neuron connectivity and odor selectivity influences the function of the piriform network. We developed a computational model consisting of a population of inhibitory neurons whose number of outgoing connections onto excitatory neurons spans a wide range (**Fig. 4a**). We compared the function of two networks: a naïve network with randomly initialized connectivity and an experienced network in which inhibitory neurons that were more differentially responsive to a set of “familiar” stimuli had a greater number of outgoing connections. Simulations of the experienced network revealed that the responses of excitatory neurons to the familiar stimuli were more discriminable than their responses to “novel” stimuli (**Fig. 4b, Extended Data Fig. 10a**). This effect was attributable to inhibitory currents onto excitatory neurons having higher variance across the set of familiar odors compared to the variance across the novel odors (**Extended Data Fig. 10b**). Larger current variance across a set of odors reduced the chance of two odors eliciting correlated responses, thereby increasing the dimensionality of the model excitatory neuron responses (**Fig. 4c**). Higher dimensionality in turn increased the discriminability of different odorant stimuli^47^. None of these effects were observed in the naïve network (**Fig. 4c**, **Extended Data Fig. 10a,b**).

**Figure 4.**
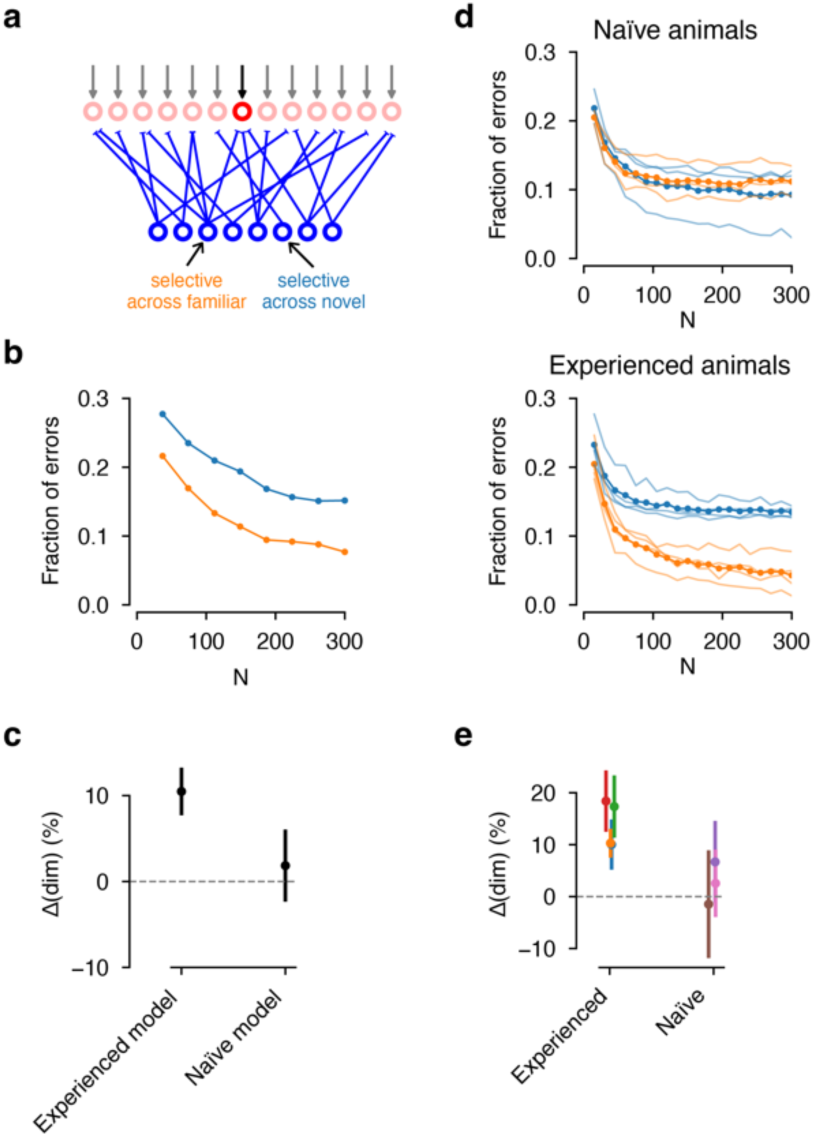
Effect of inhibitory wiring on network function. **a**, Architecture of the computational model, with inhibitory neurons (blue) projecting to excitatory neurons (red). In the experienced model, inhibitory neurons selective to familiar odors form more connections than neurons selective to novel odorant stimuli. **b**, Fraction of errors of a Hebbian linear classifier trained to perform odor discrimination based on the responses of excitatory neurons in the model (see **Methods**), plotted against the number of neurons considered. The curves are averaged across 100 model realizations, 10 cross-validation folds and separately across novel (blue) and familiar (orange) odorant stimuli. **c**, Percent difference in dimensionality between the familiar and novel odor responses in the experienced and naïve models (see **Methods**). Error bars represent 95% confidence intervals across 100 model realizations. **d**, Same as **b**, but when the classifier is trained on the responses of E neurons in our recordings. Top: naïve cohort; bottom: experienced cohort. For each value of N, the fraction of errors is averaged across 200 random E neuron subsamples. Light curves indicate individual animals, dark lines are averages across animals. **e**, Same as **c**, but for the dimensionality of the E neuron odor responses in the data. Each marker indicates a different animal, with error bars indicating 95% confidence intervals computed across different random subsampling of 500 E neurons for each dataset.

We asked whether the phenomena predicted by the model are observed in our recordings. We estimated the variance of odor-evoked inhibitory currents onto E neurons based on our measurements of the tuning of presynaptic I neurons and of the connectivity of I to E neurons (see **Methods**). Consistent with model predictions, in experienced animals the estimated variance of inhibitory currents was higher across familiar than across novel stimuli (**Extended Data Fig. 10d**), and the dimensionality of population responses was higher for familiar than for novel stimuli (**Fig. 4e**). Accordingly, piriform responses to familiar odors were more discriminable in each experienced animal (**Fig. 4d, bottom, Extended Data Fig. 10c, top**). Moreover, discriminability covaried with the amount of experience **(Extended Data Fig. 10e)**. None of these effects were observed in the naïve cohort (**Fig. 4d, top,** and **e; Extended Data Fig. 10c, bottom,** and **d, left**). Together, these results indicate that the connectivity we measured is sufficient to drive the effects observed in the model, consistent with the hypothesis that feedback inhibition decorrelates responses to odors^37^. This suggests that the experience-dependent rewiring of I neurons we have observed leads to improved discriminability of odors that the animal has encountered in its recent history.

## Discussion

We identified monosynaptically connected pairs of neurons in the piriform cortex and examined the relationship between connectivity and odor-evoked neural activity. The sparseness of the piriform network minimizes the confounds that have historically hindered efforts to reliably infer monosynaptic connectivity from spikes^21,24–26^. However, such sparse connectivity required that we obtain hundreds of simultaneously recorded neurons in a single network. This was made possible by the use of a next-generation silicon probe^11^ and the cytoarchitecture of the anterior piriform cortex, which yielded millions of recorded pairs.

Our analysis of connected pairs revealed that neurons with similar tuning to novel odors were more likely to be connected, raising the question of how this like-to-like connectivity emerges. Experience has been hypothesized to increase connections between similarly tuned neurons^42–44^. However, we found no evidence that repeated encounters with odors enhance like-to-like connectivity in the piriform. In the visual cortex, like-to-like connectivity is refined by visual experience during development^43,48^. It therefore remains possible that like-to-like connectivity in the piriform is established during a critical period during development. Alternatively, this like-to-like connectivity motif may not arise from experience and instead may reflect predetermined network constraints^49^ that we are not observing.

Experience nonetheless leaves a lasting effect on the piriform network. We discovered that the numbers of inputs to, and outputs from, inhibitory interneurons scale with how differentially they respond to odorant stimuli in the animal’s environment. How does this relationship between interneuron selectivity and connectivity emerge in the piriform network upon odor experience?

In one model, the subset of inhibitory neurons that happen to exhibit differential responses to recently encountered odors increases their numbers of inputs and outputs. This would imply the existence of an experience-dependent plasticity mechanism that depends only on the inhibitory neuron’s stimulus selectivity, rather than on the relationship between the activity of the inhibitory neuron and that of its excitatory pre- or postsynaptic partners. Such a cell-intrinsic, non-Hebbian learning rule would produce a network in which highly selective inhibitory neurons are more densely connected than other interneurons. These interneurons would need to maintain an estimate of how differentially they respond to recent stimuli. Differential responses would drive the formation or strengthening of both incoming and outgoing synaptic connections^50,51^, while uniform responses would result in a reduction in connectivity. We speculate that a mechanism for such a rule could include a factor whose concentration depends on this variance in the inhibitory neuron’s responses across stimuli, and whose presence determines the growth and retraction of axon and dendrite.

In an alternate model, the number of inputs and outputs of inhibitory neurons do not vary with odor experience. The reorganization we have observed may instead reflect changes in the selectivity of inhibitory interneurons rather than changes in their connectivity. Under this model experience results in the development of differential responses to familiar odors among those inhibitory neurons that are already highly interconnected. However, I neuron selectivity was no different between naïve and experienced animals and so any increases in selectivity among some neurons would have to be counterbalanced by proportional reductions in the selectivity of others. Ultimately, distinguishing between these two alternatives will require longitudinal observation of synaptic connectivity over the course of experience.

Experience enhances an organism’s ability to discriminate the sensory features of its environment^1–3^. Prior reports have shown that stimulus responses decorrelate with experience^52–59^. However, Hebbian plasticity, which is based on positive correlations between pre- and post-synaptic excitatory neuron firing rates, tends to correlate a network’s responses^60,61^, thus reducing discriminability. We find evidence of a process that produces the opposite effect: decorrelation and improved discriminability of commonly encountered stimuli. An experience-dependent reorganization of the network, in which the most selective inhibitory interneurons are also the most densely connected, may confer to the organism the capacity to better discriminate, recognize, and respond to the features of its environment.

## Methods Summary

Procedures performed in this study were conducted according to US National Institutes of Health guidelines for animal research and were approved by the Institutional Animal Care and Use Committee of Columbia University. See **Full Methods**.

## Data availability

Data will be made available upon reasonable request to the corresponding authors.

## Code availability

Code will be made available upon reasonable request to the corresponding authors.

## Acknowledgements

We thank L. F. Abbott, D. Aronov, C. Kentros, M. A. Long, S. McKenzie, and K. Svoboda for comments on the manuscript; G. W. Johnson and T. Tabachnik for assistance with instrumentation; F. Collman, J. Colonell, B.P. Danskin, T.D. Harris, N.A. Steinmetz, and M.A. Triplett for advice; G. Buzsáki and members of his laboratory, C. Clopath, B. Doiron, R.W. Friedrich, J.V. Georgieva, W. Gerstner, K.D. Harris, T. O’Leary, A.M.M. Oswald, D. Rinberg and members of his laboratory, and S. Sadeh for comments; M. Gutierrez, C. H. Eccard, and A. Nemes for general laboratory support; and the Howard Hughes Medical Institute for financial support. S.P.M. and A.L.-K. were supported by the Kavli Foundation, the Gatsby Charitable Foundation, the Burroughs Wellcome Foundation, and the N.I.H. award R01EB029858. S.P.M. was also supported by the Simons Collaboration on the Global Brain.

## Author contributions

This work is the result of a close collaboration between C.E.S. and A.J.P.F., who conceived the study. The experiments were performed by A.J.P.F., M.I.H., and C.E.S. The connectivity inference method was developed by A.J.P.F., S.P.M, S.W., A.L-K, and C.E.S. The computational model was developed by S.P.M. and A.L-K. The ground-truth dataset was provided by D.F.E. The data were analyzed and the manuscript was written by A.J.P.F., S.P.M., R.A., A.L-K, and C.E.S.

---

**Extended Data Figure 1.**
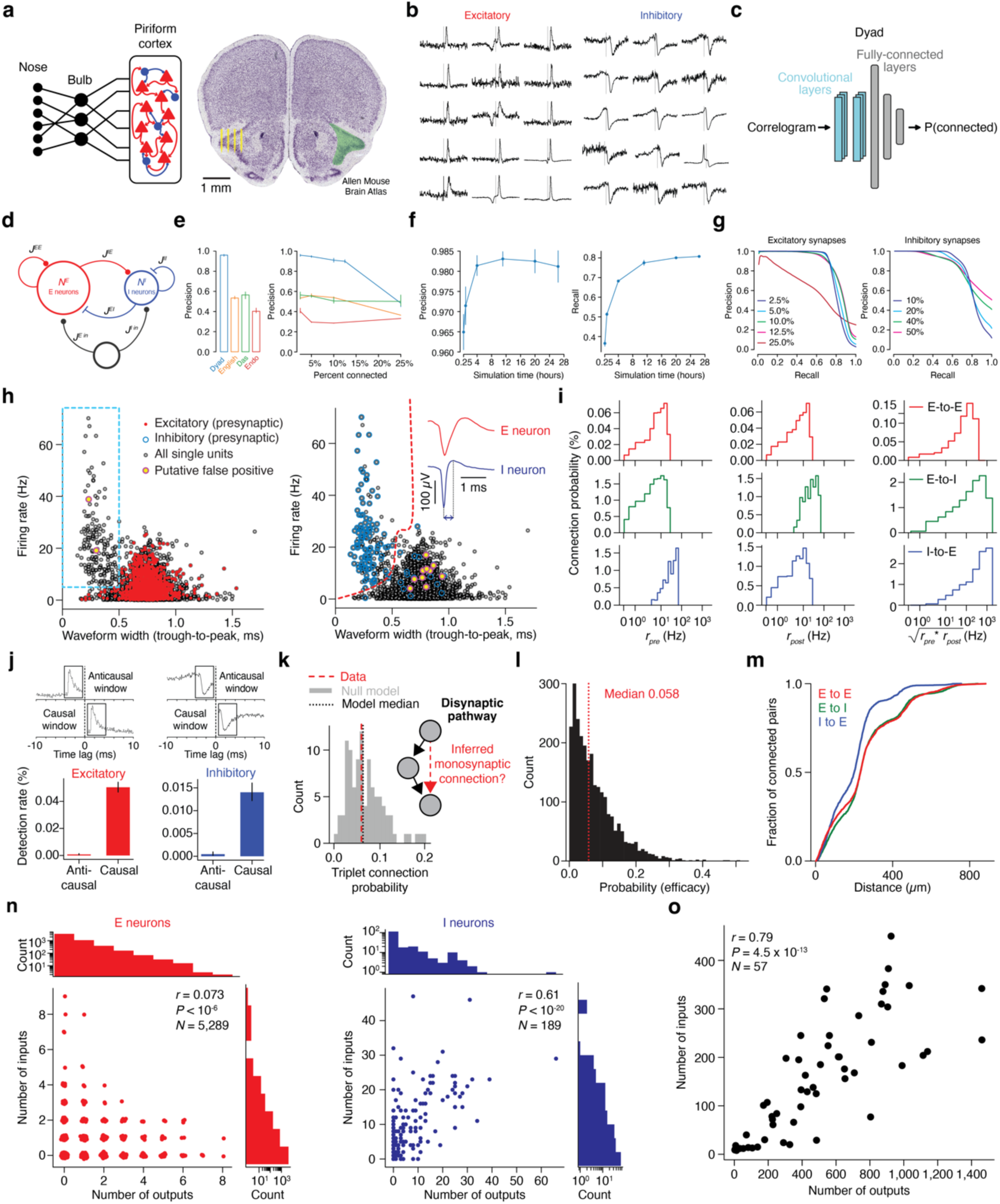
Validation of synapse inference method. **a**, Top: Simplified diagram illustrating the architecture of the mouse olfactory system. Bottom: Nissl stain from Allen Brain Atlas. Yellow marks, probe shanks drawn to scale. Green shading, anterior piriform cortex. The cell dense layer 2 and layer 3 span the four shanks of a Neuropixels 2.0 probe. **b**, Representative examples of CCGs corresponding to pairs identified by Dyad as being monosynaptically connected via an excitatory (left) or inhibitory (right) synapse. **c,** Dyad architecture. Dyad consists of two convolutional layers followed by three fully-connected layers and it was trained to classify pairs as connected based on their correlogram (see **Methods)**. The convolutional layers enabled Dyad to learn the distinctive features of cross correlograms of monosynaptically connected pairs, while maintaining flexibility, within a narrow window, with respect to the exact latency of the peak that characterizes such correlograms. **d – g**, Validation of Dyad on Synthetic ground-truth data. **d**, Illustration of the model used to generate synthetic data (see **Methods**). **e,** Left: Precision of Dyad compared to previously-published methods in detecting monosynaptically connected pairs in simulated ground truth data (see **Methods** and full precision-recall curves in Fig. 1d-bottom), when the detection threshold is set to obtain 70% recall. Right: Precision at 70% recall for different values of connectivity density of the simulated ground truth network. The left and center panel show results obtained for a simulated network with 2.5% connection probability. Previously published methods: English^22^ (peak-detection algorithm); Das^21^ (GLM-based inference); Endo^23^ (deep learning approach). **f**, Precision (left) and recall (right) at a fixed threshold, as a function of simulation time in a simulated network with 2.5% connection probability. For all the other panels, 14,400 seconds of simulated time were used. **g**, Precision-recall curves for excitatory (left) and inhibitory (right) connections, for various levels of connectivity density, averaged over three network realizations and three random subsampling of the single units. The connection probability in the legend indicates the connection probability of excitatory (left) or inhibitory (right) connections. **h**, Identification of E and I neurons. For all recorded single units, we plot their firing rate across the whole session, against their waveform width (trough-to-peak, see inset). The red dashed line in the right panel indicates the boundary used to classify single units as E or I (see **Methods**): single units that lie to the left of it are classified as I and those that lie below are classified as E. This boundary is defined by a nonlinear SVM trained using the labels shown in the right and bottom panels (see **Methods)**. The colored circles overlaid indicate whether at least one excitatory (left, red circles) or inhibitory (right, blue circles) connection was found outgoing of that single unit. Points were jittered along the x-axis to ease visualization. The dashed cyan box illustrates the definition of I neurons used to estimate Dyad’s precision in inferring excitatory synapses: firing rate > 5 spikes per second, trough-to-peak < 0.5 ms (see **Methods**). Yellow-filled circles indicate putative false positives (see **Methods**). **i**, Dependance of connection probability on firing rate, separately for E-to-E, E-to-I, and I-to-E neuron pairs. The firing rate was calculated over the course of the entire session, either for the presynaptic neuron of the pair (left column), the postsynaptic neuron (right column), or the geometric mean of the two (right column). **j**, Assessing Dyad’s precision using CCG peaks on anticausal side (see **Methods**). Top: asymmetric peaks skewed to the right in the anti-causal window (top) are considered false positives. Bottom: Detection rate (connections found divided by the total number of pairs) of Dyad in the anticausal window compared to causal one, for excitatory (left) and inhibitory (right) connections. Dyad detected 12 excitatory and 7 inhibitory false positives out of 1,490,149 total pairs (N = 2 mice). **k**, Assessing the prevalence of disynaptic chains (see **Methods**). Red dashed line: probability of triplet motifs, in which the first single unit of a disynaptic pathway also connects to the last; gray histogram: the same connection probability for a null model in which connections are shuffled while preserving the in- and out degrees of individual single units; blue dashed line: median of the null model. **l**, Distribution of efficacies for all excitatory synapses detected by Dyad. The efficacy is defined as the probability of the postsynaptic single unit emitting a spike between 0.5 and 3 milliseconds after the presynaptic spike. Red dashed line: median. **m**, Cumulative fraction of connected pairs, against the distance between the single units in the pair. **n**, Left: Number detected excitatory inputs to, and excitatory outputs from, E neurons. In this plot the points were jittered to ease the visualization. Right: incoming excitatory connections against detected outgoing inhibitory connections in I neurons. *r*, P: Pearson’s correlation coefficient and corresponding P-value. Top, distributions of the number of excitatory inputs to (right), and inhibitory outputs from (top) these I neurons. **o**, Connection counts for full-cell reconstructions of neocortical (inhibitory) basket cells imaged under large-scale serial electron microscopy^29,31^. The number of excitatory inputs onto a basket cell is strongly correlated with its number of inhibitory outputs onto excitatory neurons (Pearson’s *r* = 0.79, P < 10^-10^, N = 57 neurons from one mouse). *r*, P: Pearson’s correlation coefficient and corresponding P-value.

**Extended Data Figure 2.**
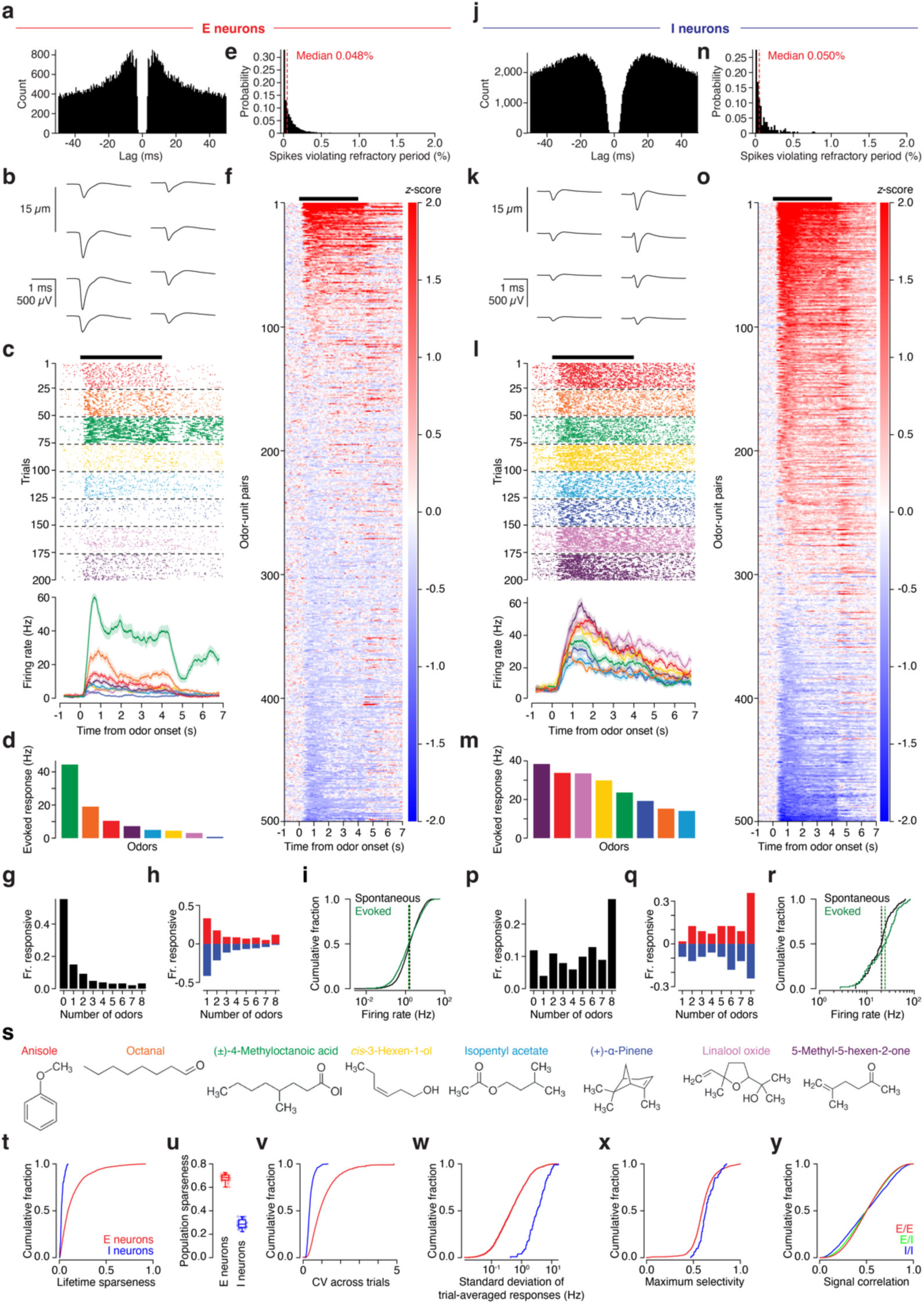
Odor-evoked response properties of E and I neurons in the piriform a–d, Response properties of an example E neuron. **a**, Spike time autocorrelogram. **b**, Mean action potential waveforms recorded on the 8 electrode sites of the probe that detected the highest-amplitude signals. **c**, Top: spike raster of odor-evoked responses, aligned to the valve opening time. Trials are sorted by odorant stimulus then trial number, and spikes (markers) are colored according to odorant stimulus. Bottom: Peristimulus time histograms, same color scheme as above. **d**, Trial-averaged, baseline-subtracted evoked firing rates in a two-second window following stimulus onset, sorted in descending order. **e**, Distribution of the percentage of spikes violating the refractory period (inter-spike interval < 1.5 msec, see **Methods**) across the population of E neurons. **f**, Heatmap showing z-scored trial-averaged stimulus-evoked responses of 500 randomly-selected E neuron-odor pairs, sorted from high increase in evoked rate (red) to high decrease in rate (blue). The z-score was computed based on a 2-sec window preceding stimulus onset to estimate the mean and standard deviation. The black bar indicates the time window in which odors are presented. **g**, Fraction of E neurons responsive to a given number of odor stimuli. Responsiveness was assessed using a paired Wilcoxon signed-rank test between the number of spikes in the two seconds preceding and following odor presentation, with a threshold of 10^-4^ on the P-value. **h**, Same as **g**, but separating neurons whose activity increased after odor delivery (red) from those whose activity decreased (blue). **i**, Cumulative distribution of spontaneous (black) and stimulus-evoked (green) firing rates; medians indicated in dashed lines. The stimulus-evoked rate was computed using a 2-sec window following stimulus onset. **j–m**, Same as **a–d**, but for an example I neuron. **n–r**, Same as **e–i**, but for the population of I neurons. **s**, Odorants employed and their molecular structure. **t**, Cumulative distributions of lifetime sparseness, for E and I neurons. In panels **t–y**, red and blue correspond to E and I neurons, respectively. See **Methods** for details about how each statistic was computed. **u**, Population sparseness. The box indicates the 1^st^ and 3^rd^ quartile (horizontal line: median), and the whiskers indicate the full range of the data. Each marker corresponds to one odor stimulus and one animal. **v**, Cumulative distributions of the coefficient of variation of single-neuron responses across trials. **w**, Cumulative distributions of the standard deviation of trial-averaged responses. **x**, Cumulative distributions of maximum selectivities. For each neuron-odor pair, the selectivity was defined as the average one-vs-one classification performance on a trial-by-trial basis. The maximum selectivity was obtained for each neuron by taking the maximum across odors. **y**, Cumulative distributions of signal correlation, for E/E, E/I, and I/I pairs.

**Extended Data Figure 3.**
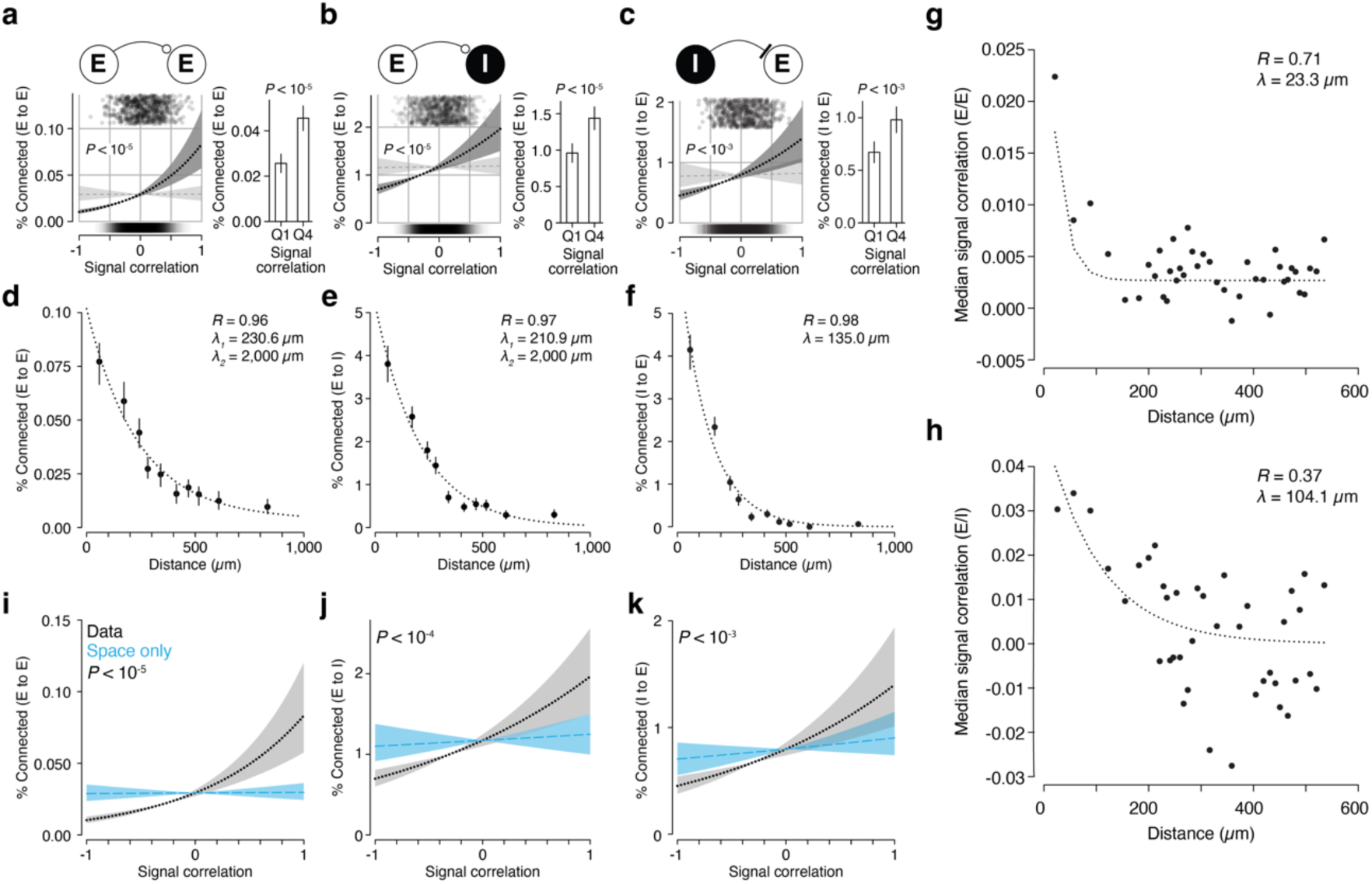
Like-to-like connectivity and spatial structure in the piriform cortex. **a–c**, Connection probability as a function of signal correlation, as in Fig. 2b, separately for **a**, (E-to-E, reproduced from Fig. 2b to permit comparison), **b**, E-to-I, and **c**, I-to-E. **d–f**, Connection probability as a function of distance between the pair, separately for E-to-E (**d**), E-to-I (**e**) and I-to-E (**f**) connections. Pairs were divided into 10 distance bins so that each bin included the same number of pairs. Markers: connection probability in each distance bin, error bars: 95% confidence intervals obtained via bootstrapping; dashed lines: double exponential fit to the binned data. To assess whether the spatial structure we found is consistent with previous findings, we fixed the spatial scale of one exponential to 2 mm, as reported in prior measurements over longer distances^8,9^. For I to E connections, the fit returned zero for the coefficient corresponding to the 2mm-wide exponential, indicating that the spatial is best fit by a single exponential. This was not the case for E to E and E to I connections, indicating that our findings are compatible with previous reports, and capture an additional, fast decay component that was not resolvable by the methodology in previous reports. **g,h**, Signal correlation as a function of distance between the pair, separately for E/E (**g**) and E/I (**h**) pairs. Pairs were separated in 40 equally spaced bins and for each bin the median across pairs of the signal correlation was measured. Dashed lines indicate an exponential fit to the binned data, with spatial scale reported as λ in the figure. *r* and corresponding P-values indicate Pearson’s correlation coefficients of the non-binned data. **i–k,** Connection probability as a function of signal correlation for a null model that preserves the spatial structure of the piriform (see **Methods**), for E-to-E (**i**), E-to-I (**j**), and I-to-E (**k**) connections. Blue dashed lines and shading indicate logistic fits and 5th and 95th percentile of the fit to such null model. Black dotted lines and shading are reproduced from **a-c** to permit comparison.

**Extended Data Figure 4.**
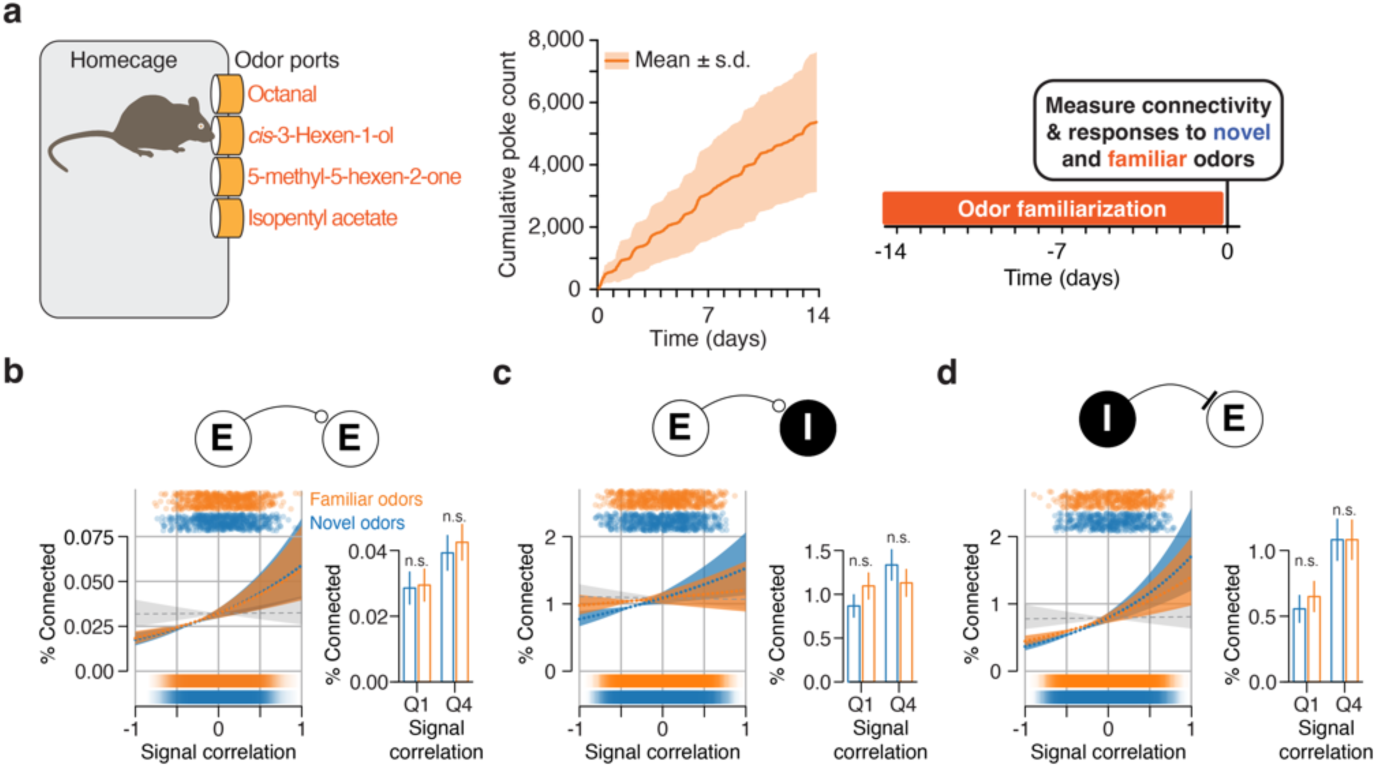
Experience does not enhance like-to-like connectivity in the piriform cortex for E-to-E, E-to-I, or I-to-E pairs. **a**, Left: Diagram of the apparatus for familiarization with odors and the four odorants used in that protocol. Center: Cumulative number of times that a mouse sampled the odor ports, averaged across N = 4 mice. Right: experiment timeline. Animals underwent this familiarization protocol for approximately two weeks (orange bar) followed by a single recording session to measure synaptic connectivity and odor responsiveness. During this recording the animals were presented both the four familiar odorants they had sampled in their home cage as well as four novel stimuli. **b-d**. Connection probability as a function of signal correlation across novel or familiar odorant stimuli separately for **b**, E-to-E (reproduced from Fig. 2d to permit comparison), **c**, E-to-I, and **d**, I-to-E.

**Extended Data Figure 5.**
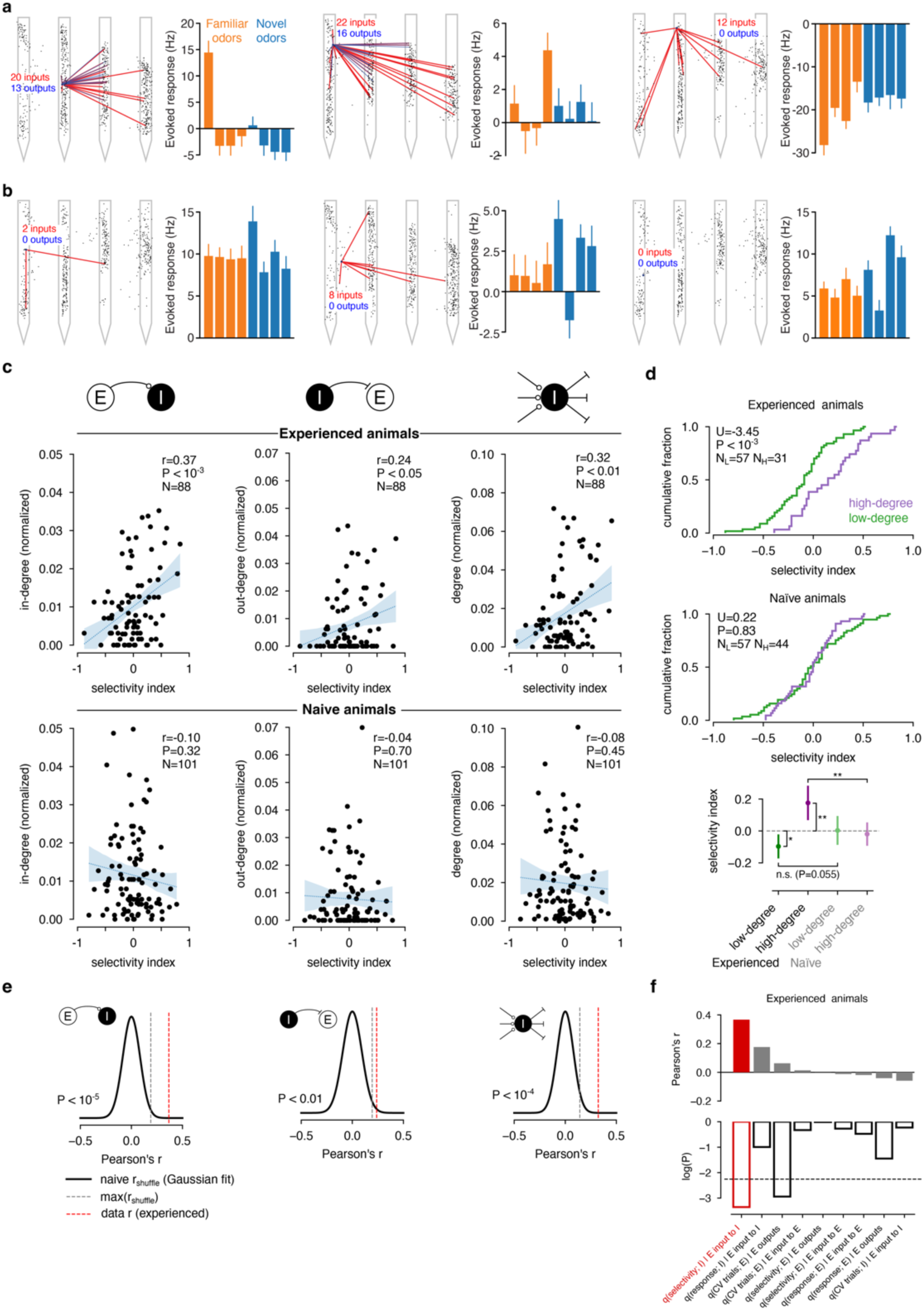
The effect of experience on the relationship between the connectivity and selectivity of I neurons. **a**, Connectivity maps (left) and evoked response amplitudes (right) of the three I neurons with the highest positive selectivity index, after the example shown in Fig. 3. Black circles: estimated single unit locations; red lines: incoming excitatory connections; blue lines: outgoing inhibitory connections; grey masks: the four silicon probe shanks. **b**, Same as **a**, but for the three I neurons with the highest negative selectivity index. **c**, Top row: In degree (left), out degree (center), and degree (right) of I neurons in experienced animals as a function of their selectivity; degrees are normalized by the number of single units in each recording to permit pooling of multiple datasets. A single unit’s selectivity was computed using a standard difference index (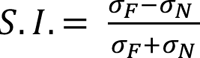, where σ_*N*_, σ_*F*_ are the standard deviation of the trial-averaged odor response across novel or familiar odors, see **Methods**). Statistics report Pearson’s correlation coefficients and corresponding P-values with respect to a null distribution in which the single unit’s selectivities and degrees were independently shuffled. The blue dotted lines show a linear fit and shaded regions indicate the 95% confidence interval of the fit. Bottom row: Same as top, but for naïve animals. Each of the dependencies we observed in experienced mice (panel **c**) differed significantly from those found in naïve animals (panel **e**): in-degree: t = 3.13 P = 1.0 x 10^-6^, N = 101, 88 I neurons (experienced, naïve), out-degree: t = 1.8, P = 1.15 x 10^-3^, N = 101, 88 I neurons (experienced, naïve), degree: t = 2.71, P = 1.4 x 10^-5^, N = 101, 88 I neurons (experienced, naïve). **d**, Top: cumulative distributions of S.I.s for I neurons that are high degree (> 0.015, purple), and low degree (< 0.015, green), for experienced (top) and naïve (middle) animals. Statistics result from a Wilcoxon rank-sum test. Bottom: mean S.I. and 95% confidence intervals (assuming normality) for high- and low-degree I neurons, for experienced and naïve animals. Significance was tested using the T-test (when comparing means to zero) and the Wilcoxon rank-sum test (when comparing two different single unit groups). Experienced, high degree vs zero: P = 0.0037; t = 3.14, N = 31 I neurons; low degree vs zero: P = 0.016, t = −2.50, N = 57 I neurons. High degree, experienced vs naïve: P = 0.0069; U = −0.52, N = 31, 44 I neurons (experienced, naïve). Low degree, experienced vs naïve: P = 0.055, U = 0.07, N = 57, 57 I neurons (experienced, naïve). **e**, Distribution of Pearson’s *r* correlation coefficients between SI and in degree (top), out degree (middle) and degree (bottom) obtained when considering all 70 ways to split the eight odorant stimuli in a set of “novel” and a set of “familiar” odors, in naïve mice. The black lines show Gaussian fits to these distributions, the gray dashed lines indicate the maximum *r* across all possible partitions, and the red dashed line indicates the *r* value obtained in experienced mice (same value as shown in **c**-top). **f**, Pearson’s *r* correlation coefficients (top) and corresponding P-values (bottom) between the degree and difference index *q(x)* of single-neuron response properties in experienced animals. 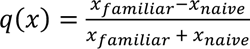, where *x* can be the trial-averaged response (“response”), the standard deviation across odors of the trial-averaged response (“selectivity”), or the coefficient of variation across trials (“CV”). Bars in both panels are sorted according to *r*. The black dashed line indicates the threshold for significance after Bonferroni correction for 9 comparisons (α = 0.0055).

**Extended Data Figure 6.**
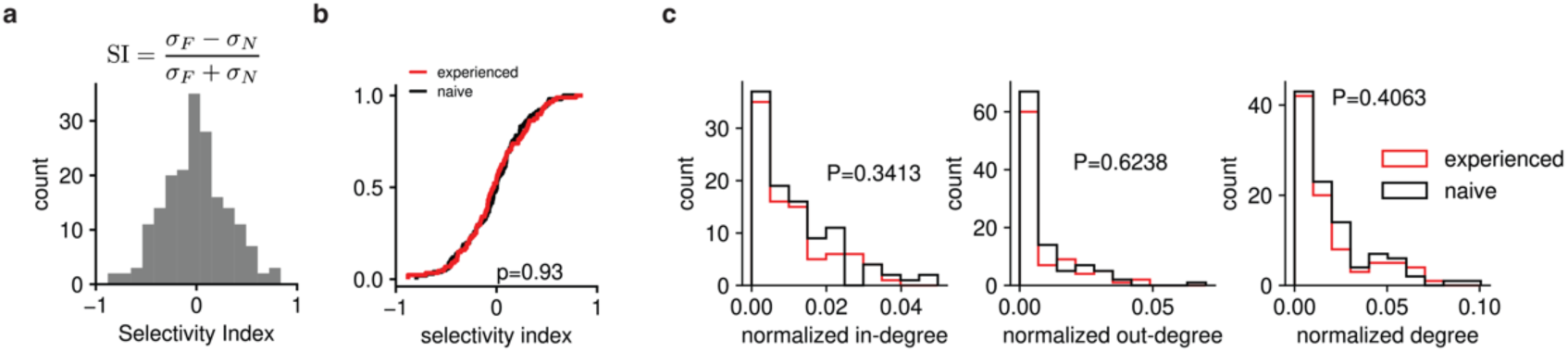
Distributions of odor selectivity and degree are unaffected by experience. **a**, Distribution of SIs across all I neurons in all animals. **b**, Cumulative distributions of S.I.s for I neurons in experienced (red) and naïve (black) animals. The P-value was computed using the Kolmogorov-Smirnov test (N_naïve_ = 101; N_experienced_ = 88). **c**, Distributions of I neuron in degrees (left), out degrees (center), and degrees (right), each normalized by the number of single units in each recording to permit pooling across recordings. The distributions are plotted separately for naïve (black) and experienced (red) mice, and P-values are computed using the Kolmogorov-Smirnov test (N_naïve_ = 101; N_experienced_ = 88).

**Extended Data Figure 7.**
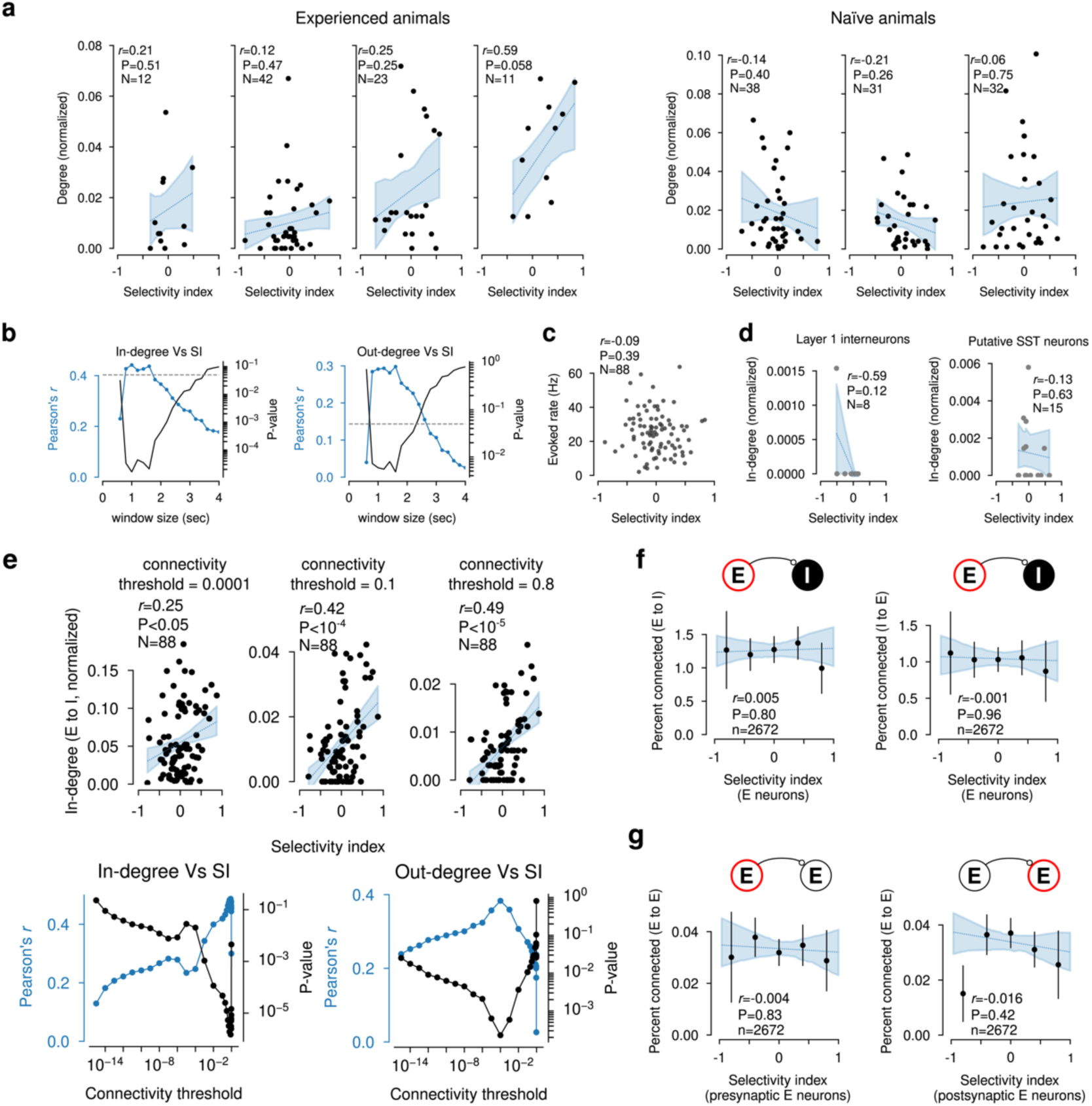
Robustness of this study’s principal findings. **a**, Degree as a function of S.I., separately for each mouse, as in **Extend Data Figure. 5c. b**, Effect of varying the size of the quantification window used to compute odor responses. Left: Pearson’s correlation coefficient (blue) and corresponding P-value (black) between I neuron out degree and S.I. in experienced animals as a function of quantification window size. Grey dashed line: P = 0.05. Right: same but for in degree. In all the analyses in this study we employed a 2-second quantification window. **c**, Odor-evoked firing rate as a function of S.I. for all I neurons in experienced animals. **d**, Normalized in-degree of layer-1 interneurons and putative somatostatine-positive (SST) neurons, as a function of their selectivity index. **e,** Robustness of results with respect to the choice of the threshold employed to consider a pair connected. The only inclusion criteria was the threshold without any further manual curation. Top: The normalized in-degree of I neurons increases with the selectivity index, for three different values of the connectivity threshold. Bottom: Pearson’s *r* and corresponding P-value between selectivity index and the normalized in-degree (left) and out-degree (right), as a function of the connectivity threshold. **f, g**, Connection probability did not depend on the selectivity of E neurons. **f**, Connection probability as a function of the selectivity index of the E neuron of the pair for E-to-I (left) and I-to-E (right) neurons. **g**, E-to-E connection probability, as a function of either the pre-synaptic (left) or post-synaptic (right) E neuron’s selectivity.

**Extended Data Figure 8.**
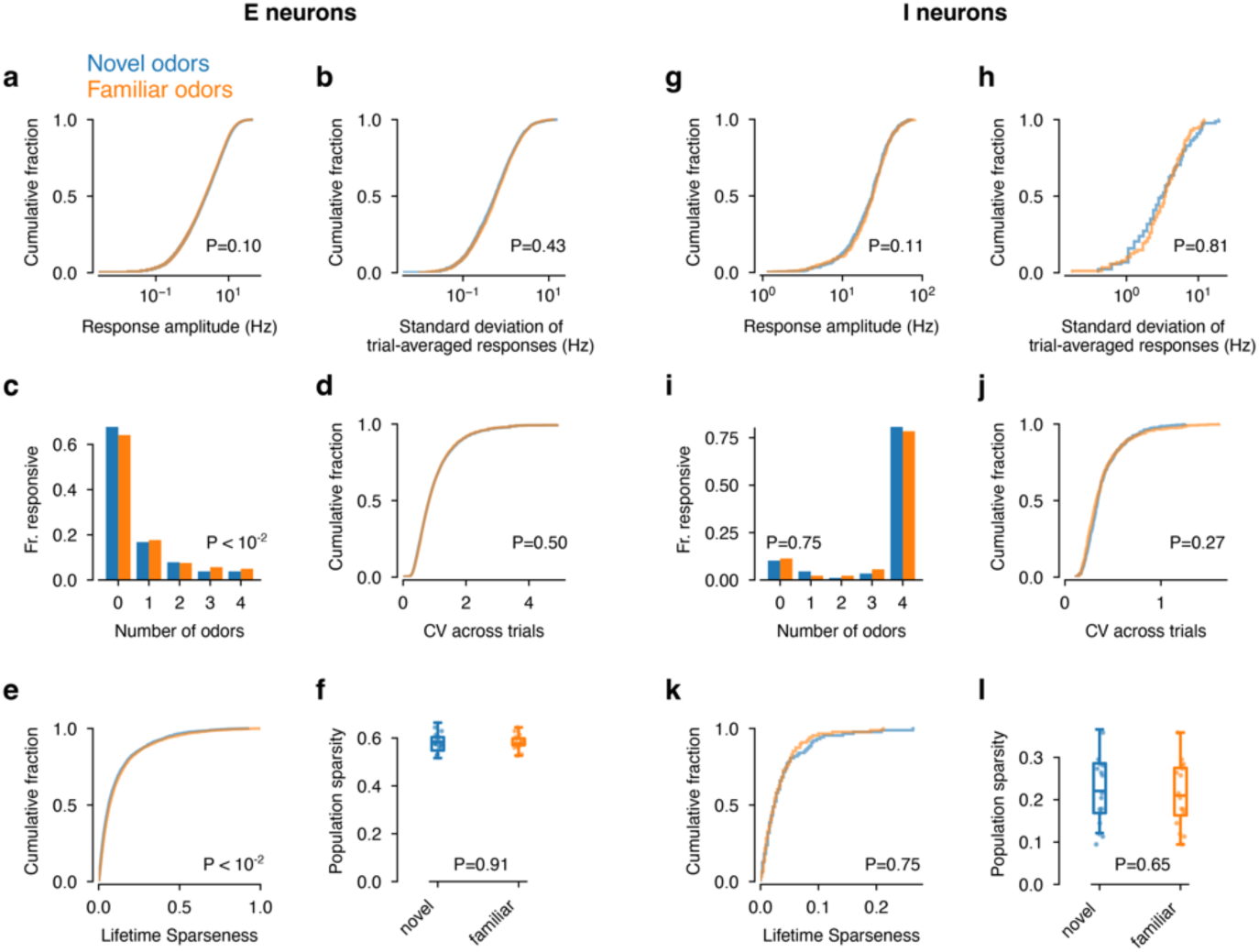
Odor-evoked response properties of E and I neurons in experienced animals Throughout this figure blue indicates novel and orange familiar odors. Unless stated otherwise, all P-values were computed using a Wilcoxon rank-sum test. See **Methods** for details about how the other statistics were computed. **a–f**, Odor-evoked responses of E neurons (N = 2,672). **a**, Cumulative distributions of trial-averaged evoked firing rates. **b**, Cumulative distributions of the standard deviation of trial-averaged evoked firing rates. **c**, Fraction of E neurons responsive to a given number of odors in the novel or familiar set. Responsiveness was quantified by comparing the number of spikes in the two-second window after valve opening to the number of spikes in the two-second window before valve opening. A single unit was considered responsive to an odor if the corresponding P-value was smaller than 10^-4^. The difference between the two distributions was assessed using the Mann–Whitney *U* test. Although the distribution is only marginally higher for familiar odors, the effect was significant (P = 0.00182). **d**, Cumulative distributions of the coefficient of variation (CV) of individual E neurons across trials. **e**, Cumulative distributions of lifetime sparseness of individual E neurons. Although the lifetime sparseness was only marginally larger for familiar odors, the effect was significant (P = 0.00178). **f**, Population sparseness (see **Methods**) across the population of E neurons. The box indicates the 1^st^ and 3^rd^ quartile (horizontal line: median), and the whiskers indicate the full support of the distribution of sparsities across odors and animals. **g-l,** Same as **a-f**, but for the odor-evoked responses of I neurons (N = 88).

**Extended Data Figure 9.**
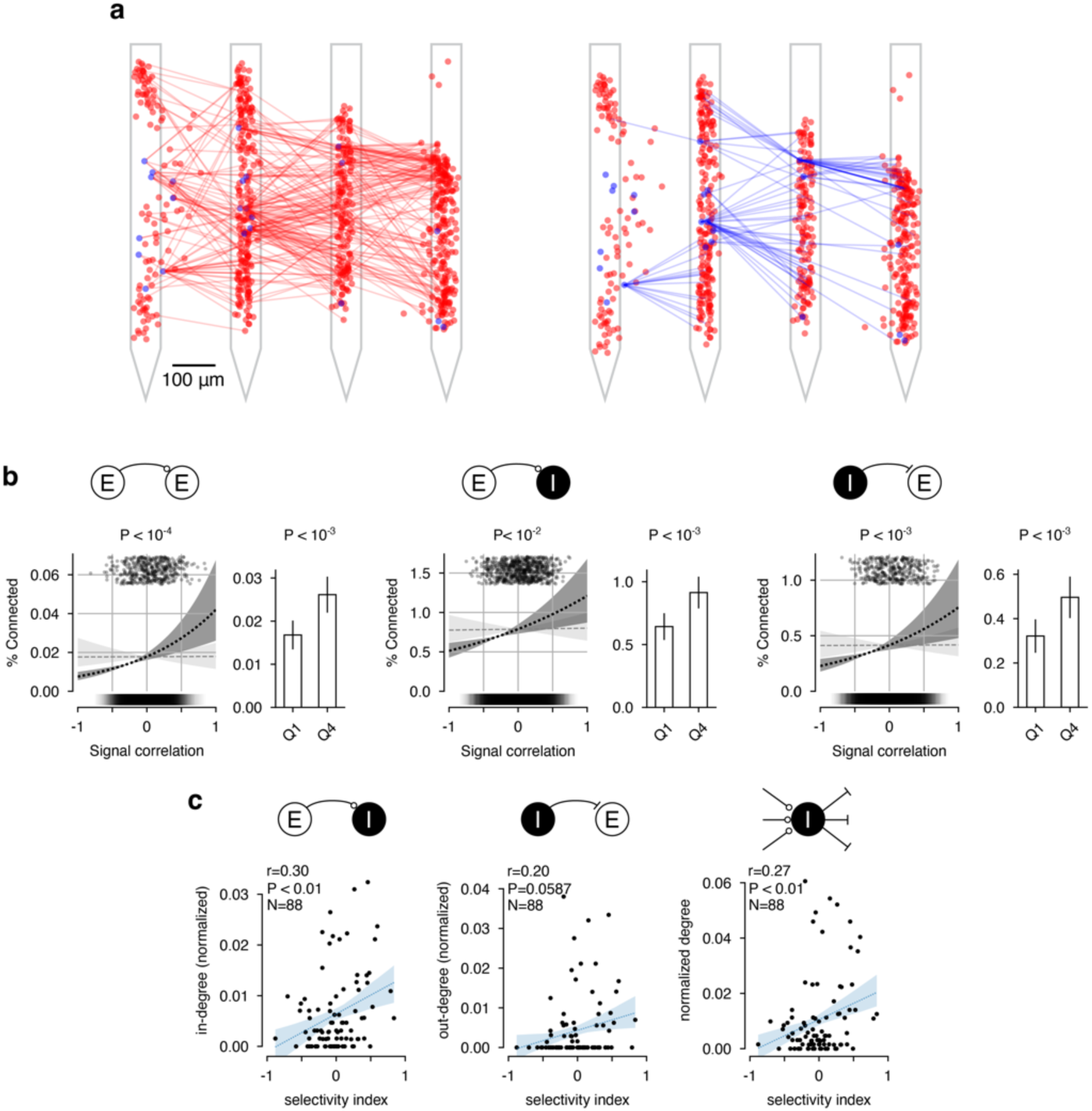
Principal findings hold when considering only cross-shank connections. **a,** Example of excitatory (left) and inhibitory (right) connectivity for one dataset, as in Fig. 1e but considering only connections across different probe shanks, which are separated by 250 µm center-to-center. **b,** Connection probability as a function of signal correlations for E-to-E (left), E-to-I (center) and I-to-E (right) pairs, when considering only cross-shank connections. Plots and quantifications are as in Fig 2b and **Extended Data** Fig. 3a–c. **c,** Normalized in degree (left), out degree (center), and degree (right) for I neurons in experienced animals as a function of their selectivity index, when considering only cross-shank connections. Plots and quantifications are as in **Extended Data** Fig. 5.

**Extended Data Figure 10.**
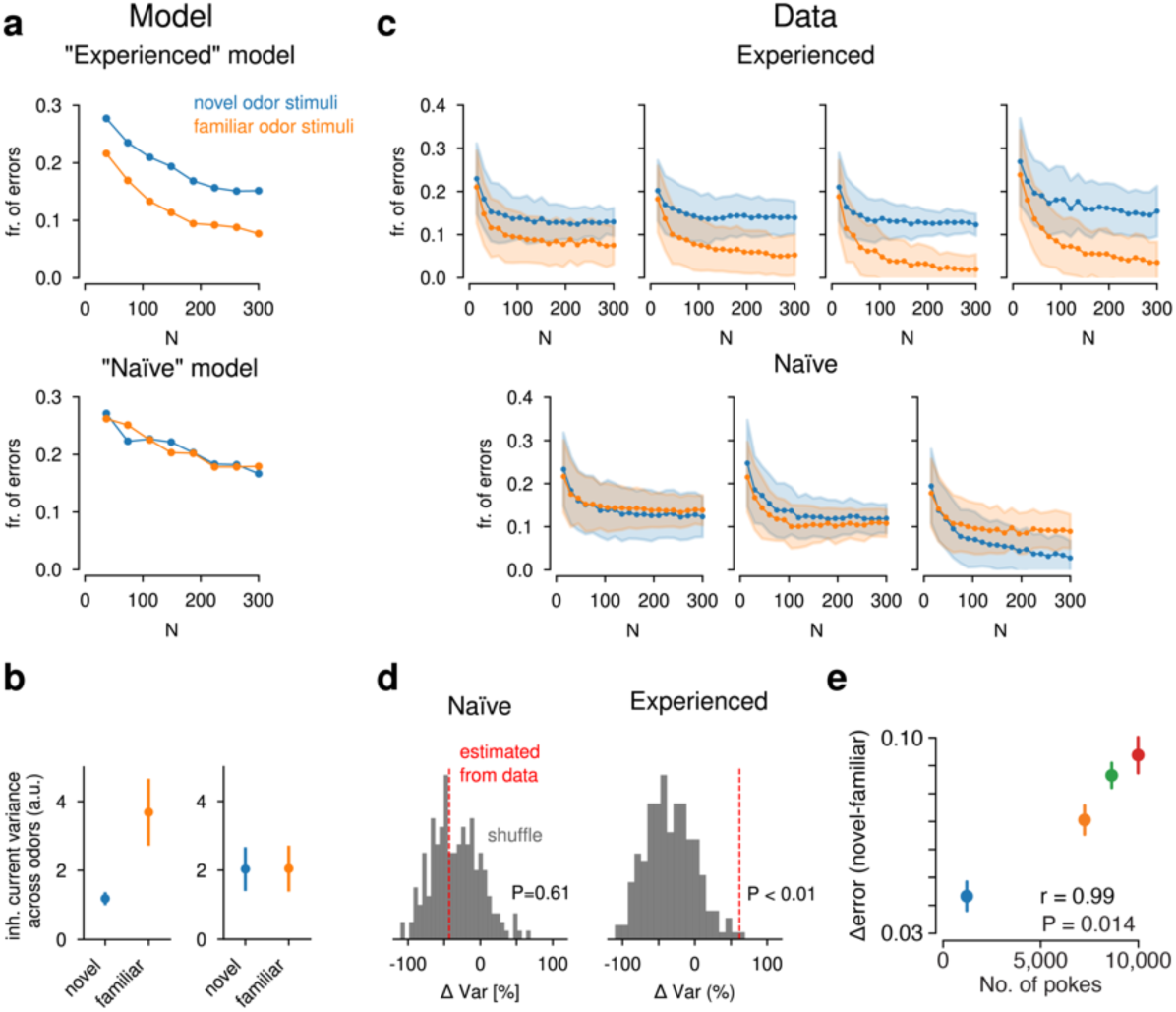
Increased variance of inhibitory input onto E neurons decorrelates their responses. **a**, Fraction of error of a Hebbian linear classifier, as in Fig. 4a, but showing the results for both the experienced (top, reproduced from Fig 4a to permit comparison) and the naïve (bottom) model. **b,** Variance across odors of the total inhibitory current onto excitatory neurons in the model, for both the experienced (left) and naïve (right) model. The variance is computed separately across novel and familiar odors, and averaged across 100 model realizations. Error bars: 95% confidence intervals for the mean variance. **c**, Fraction of errors of a Hebbian classifier trained on the odor responses of E neurons, as in Fig. 4d, for each individual mouse for novel (blue) and familiar (orange) stimulus set. Solid curves: mean; shaded area: standard deviation across odors and subsamples. **d**, Percent change in variance estimated from our electrophysiological recordings (dashed red line, see **Methods**), for naïve (left) and experienced (right) mice. Gray histogram: percent change in variance that the same estimation yields when connectivity is shuffled. The reported P-values are obtained with respect to this null model. **e,** Difference in classification performance between novel and familiar odorant stimuli (differences from panel **c**, averaged across N), as a function of the amount of experience (number of nose port pokes) the mouse had with the set of odors (Pearson’s *r* = 0.99, P = 0.014, N = 4 mice).

## Methods

### Ethical compliance

All procedures were approved by the Columbia University Institutional Animal Care and Use Committee (protocol AC-AABH6557) and were performed in compliance with the ethical regulations of Columbia University as well as the Guide for Animal Care and Use of Laboratory Animals.

### Animals

We employed 7 male C57BL/6J adult mice (Jackson laboratories, Bar Harbor, ME) aged 15 to 34 weeks (26.0 ± 9.6 weeks, mean ± s.d.). Animals had free access to food and water and were acclimated to the reversed light/dark cycle for at least two weeks before recordings or behavioral experiments were initiated. Mice were group-housed before acquiring odor experience and housed singly thereafter.

### Stereotactic targeting and head plate attachment surgery

As previously described^62^, briefly: Animals were anesthetized with isoflurane (3% induction, 1.5-2% maintenance); carprofen (5 mg/kg) and bupivacaine (2 mg/kg) were administered for preoperative analgesia and numbing, respectively before exposing the skull. The head was aligned as previously described, and photographs of landmarks were taken to ensure subsequent reliable targeting of the anterior piriform cortex at 1,500 µm (medial-most shank) and 2,250 µm (lateral-most shank) lateral to the midline and 1,150 µm posterior to the rostral confluence of the sinus (pRCS). A titanium head plate (G. Johnson, Columbia University) was secured to the using a thin layer of cyanoacrylate adhesive (Krazy Glue, Elmer’s Products, Atlanta, GA) applied to the skull followed by adhesive luting cement (C&B-Metabond, Parkell, Inc., Edgewood, NY). Two bone screws were then applied bilaterally and secured with the luting cement above the cerebellum to serve as ground and reference electrodes. The incision was closed with sutures and mice recovered for at least 14 days before behavioral or physiological experiments.

### Craniotomy

As previously described^62^, briefly: On the day before the recording, A craniotomy and durotomy were performed using a dental drill (Osada Success 40, Osada Electric Co., Ltd., Tokyo, Japan) and a fine scalpel blade (Fine Science Tools #11), targeted to the location based on the previously photographed landmarks. The animal was then allowed to recover overnight. In the event of pial bleeding, edema, or other signs of damage to the brain the experiment was aborted.

### Neurophysiological recordings

As previously described^62^, briefly: Experiments were conducted in dark, sound-attenuated conditions, with bandpass filtered acoustic white noise (1,000–45,000 Hz; approximately 7 dB). The animal was administered 1.0 mL of saline and then maintained awake and head-restrained with its body in a tube designed to promote calm^63^. Recordings were performed using 4-shank Neuropixels 2.0 probes^11^ aligned to the vertical axis of travel of the micromanipulator (Patchstar, Scientifica, East Sussex, United Kingdom) using a custom 2-axis gimbal (G. Johnson, Columbia University). Following penetration, in the event of dimpling of the craniotomy or bending of one of the probe shanks, the issue was either resolved on the spot or the recording was aborted. Based on previous calibration experiments^62^, which yielded unambiguous physiological signatures to establish correct targeting, the probe was targeted to approximately 3,500 µm of depth, and then fine adjustments in the vertical axis were made based on real-time monitoring of neural signals. In the anterior-most portion of the cortex the cell-dense layer folds over itself (**Extended Data** Fig. 1a). Proper targeting was achieved when spike waveforms emanating from both the dorsal and the ventral portions were visible on the probe. Failure to reach this target resulted in the recording being aborted, as we never made more than a single penetration to maximize tissue health. The final depth from the pial surface for successfully targeted preparations was 3,772 ± 291 µm (N = 7 mice, mean ± s.d.). The probe was allowed to settle for approximately one hour, after which recording was initiated. The duration of recordings ranged from 5.3 to 7.1 hours (6.1 ± 0.9 hours, mean ± s.d., N = 7 mice). These recordings comprised three phases (Fig. 2a): two stretches of spontaneous activity lasting approximately two hours each at the beginning and at the end, to collect a sufficient number of spikes to infer synaptic connections; and a middle phase lasting approximately 3 hours during which odorant stimuli were administered, to measure odor responses.

### Odorant stimuli

During this middle phase we administered 8 distinct monomolecular odorants 25 times each in pseudorandom order. Each trial included a 4-second odorant pulse followed by a mean 60 sec inter-trial interval (ITI), drawn at random from a uniform distribution between 50 and 70 seconds. Odorant stimuli were administered as previously described^62^ using a custom built, flow-equalized olfactometer. Odorant kinetics were measured using a photoionization detector (miniPID, Aurora Scientific, Aurora, ON, Canada) sampling between the nose and the exhaust line to confirm post-hoc that stimulus delivery was well controlled.

The odorants employed were: 2% anisole (Sigma cat. no. W209708), 2% octanal (Sigma cat. no. O5608), 20% (±)-4-methyloctanoic acid (Sigma cat. no. W357502), 2% *cis*-3-hexen-1-ol (Sigma cat. no. W256307), 2% isopentyl acetate (Sigma cat. no. 306967), 4% (+)-α-pinene (Sigma cat. no. 268070), 6% linalool oxide (Sigma cat. no. 62141), 4% 5-methyl-5-hexen-2-one (Sigma cat. no. 364479), dissolved (% v/v) in 15 ml Dipropylene Glycol (DPG). The odorants had diverse functional groups and organoleptic properties, and were not known to release innate attractive or aversive behaviors. In preliminary experiments they were titrated to activate responses in comparable fractions of piriform neurons.

### Data acquisition and spike sorting

Neural signals were acquired at 30 kHz with a PXI base station (imec, Leuven, Belgium) using the SpikeGLX acquisition software (https://billkarsh.github.io/SpikeGLX/). Spike sorting was performed using the ecephys pipeline (Janelia fork: https://github.com/jenniferColonell/ecephys_spike_sorting) with Kilosort 2.5^64^. The output of Kilosort was manually curated using Phy (github.com/cortex-lab/phy). We excluded from analysis any templates corresponding to electrical noise, templates that ran down or appeared mid-recording, as well as any templates for which refractory period violations exceeded 2%. The refractory period was defined as an inter-spike interval < 1.5 msec (Median 0.048% refractory period violations, Q1=0.011%, Q3=0.12%). The percentage of templates that had zero refractory period violations was 13.7% (full distributions for E and I neurons in **Extended Data** Fig. 2e,n).

### Odor familiarization

A subset of animals acquired experience with 4 out of the 8 odorants prior to the recording session. These 4 odorants, *cis*-3-hexen-1-ol, octanal, 5-methyl-5-hexen-2-one, isopentyl acetate, were diluted in DPG at the same concentration as that used in the recordings. The animals acquired experience with the stimuli in a modified home cage as previously described^45^. Briefly, four BPod poke ports (Sanworks, Rochester, NY) were placed in the wall of the home cage, with each one delivering one of the 4 odorants conditional on the animal’s nose breaking the port’s infrared beam. The animals had *ad libitum* access to food and water and could volitionally sample the stimuli over a period of approximately two weeks. The diluted odorant solutions were bubbled with air at 0.25 L per minute, and fresh stocks were switched in every one to three days to prevent depletion. Recordings were performed immediately following removal of the mice from the apparatus.

### Inference of Synaptic Connections (Dyad)

We computed spike-time cross correlograms (CCGs) for all pairs of neurons for each recording using a time bin of 0.1ms and time lags between −20 and +20 milliseconds. CCGs were pre-processed for synapse inference as follows: first, CCGs were linearly interpolated between −0.3 and 0.3ms to remove artifacts introduced by the spike-sorting software. Then, CCGs were smoothed using a 0.5ms-wide boxcar function. Finally, CCGs were normalized by the product of the firing rate of the two neurons in the pair. A small regularization constant λ = 2 Hz^2^ was used to prevent such normalization to overly weigh pairs of low-firing-rate neurons.

We trained two deep convolutional networks (Dyad) to recognize CCGs that exhibited the characteristic features of excitatory and inhibitory connections (see below). One network was trained for excitatory and one for inhibitory connections. After preprocessing the CCGs, Dyad’s input features were constructed by considering the portion of the CCG between −10 and 10 milliseconds and performing z-scoring across time lags. Each Dyad network comprised two convolutional layers (16 channels each, kernel size: 9, stride: 3 and 1 respectively), followed by three fully-connected layers of ReLU units (512, 256, and 128 units from early to late in the network). The network readout received input only from the last layer and had a sigmoidal activation function to obtain a network output between 0 and 1. The network was trained by minimizing the binary cross-entropy loss using Adam^65^ (learning rate: 10^-5^, weight decay: 0.01) for 200,000 epochs. To deal with our imbalanced dataset (unconnected pairs are ∼1,000 times more common than connected ones), each epoch we sampled an equal number of positive and negative examples. The model that exhibited the lowest loss on the validation set was selected and its performance was assessed on the test set.

To obtain the training dataset, we used a custom peak-detection algorithm based on thresholding the derivative of the CCG, followed by manual curation, to obtain two initial dataset that comprised approximately 300 positive examples each (one for putative excitatory connections, one for putative inhibitory connection). All the other pairs in the dataset were assigned a negative label.

After training, we fed the CCGs for all the pairs to Dyad and used the value of the network readout to rank all pairs. All pairs with a score smaller than 10^-4^ were considered not connected, while those above such value were manually curated.

Manual curation was based on three criteria: 1) symmetry: sometimes Dyad assigned a high-rank to CCGs which had a sharp peak exhibiting all features of a monosynaptic connection but was symmetric. Such pairs were considered not connected in this study; 2) rise-time: pairs whose peak started to rise before a lag of 0.5 milliseconds were also considered not connected; 3) near-zero fluctuations: pairs that exhibited large fluctuations for near-zero lags (−1 to 1 ms) of magnitude comparable to the main causal peak were also considered not connected. This was to exclude cases in which spike-sorting artifacts, more common for nearby units, might distort the shape of a larger peak due to common input to resemble one resulting from monosynaptic connectivity.

Our results did not depend on the details of the manual curation and also held when using the score assigned by Dyad to classify neuron pairs without any manual curation, across a large range of score thresholds (**Extended Data** Fig. 7e).

### Validation of Dyad on Ground Truth Data

We tested Dyad using a ground truth dataset with positively identified monosynaptic connections. This dataset, previously published in English et al.^22^, was generated using a combination of extracellular silicon probe recordings and juxtacellular stimulation in the hippocampal area CA1 of the mouse brain. The methods are described in detail in the original publication. Briefly, to decouple the spiking of the presynaptic neuron from the network dynamics, the juxtacelllularly-recorded neuron was stimulated at random times, reliably evoking spikes at every stimulation. We computed *evoked* CCGs using only the first spike evoked by the stimulation for each juxtacellularly-stimulated neuron. Peaks detected in such correlograms are decoupled from the network dynamics and therefore cannot be due to common input. We applied the same method described in the original publication^22^ to these evoked CCGs to identify monosynaptic connections outgoing from juxtacellularly-stimulated neurons, obtaining 30 connected pairs and 274 unconnected pairs.

We then computed *spontaneous* CCGs for all pairs of one juxtacellularly-stimulated neuron and one neuron recorded using a silicon probe, using all spikes occurring at least 10 milliseconds after the end of the stimulation. We used Dyad to assign a score to all the spontaneous CCGs, and varied the threshold applied to the output of Dyad to generate the precision-recall curve in Fig 1d.

### Validation of Dyad on Synthetic Data

We tested Dyad on synthetic data generated by a spiking neural network model consisting of two interacting populations of excitatory and inhibitory neurons. We considered a network of LIF neurons (indexed by *i*=1, …, *N*) that receive input from other neurons in the network and from an external population of *N^Ext^*neurons. Neurons emit a spike at time *t* if the membrane potential *V*^i^ reaches the firing threshold *θ*. Between spikes, for all t > t^f^ +t^ref^, where t^f^ is the last spike time and t^ref^ is the duration of the refractory period, the voltage dynamics follows

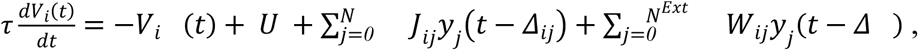

Where Δ_ij_ is the synaptic delay of from neuron *j* to neuron *i*, *U* is the resting potential, and *y*_j_(*t*) is the spike train of neuron *j* filtered using an exponential kernel with time scale τ_*syn*_. To reduce simulation time, we considered Δ_ij_ = *1*ms for our analyses. However, we also verified that introducing heterogeneous delays (Δ_ij_ = Δ_j_ sampled i.i.d. from a uniform distribution between 0.5 and 2.5 ms) does not materially alter our main results. After every spike, the membrane potential is reset to zero and held at zero for the duration of the refractory period. Neurons in the external population fire as Poisson processes at a constant rate of *A^Ext^* spikes per second. The network was simulated by discretizing time in steps of dt = 0.2ms.

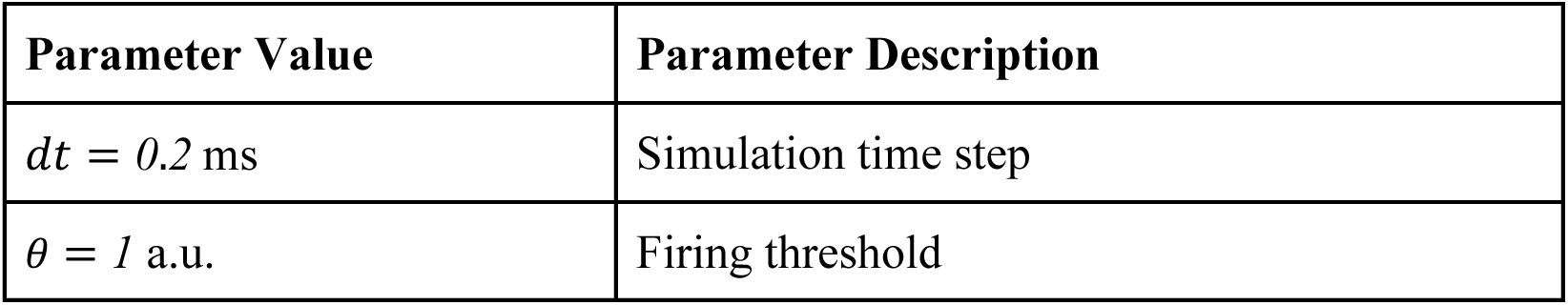
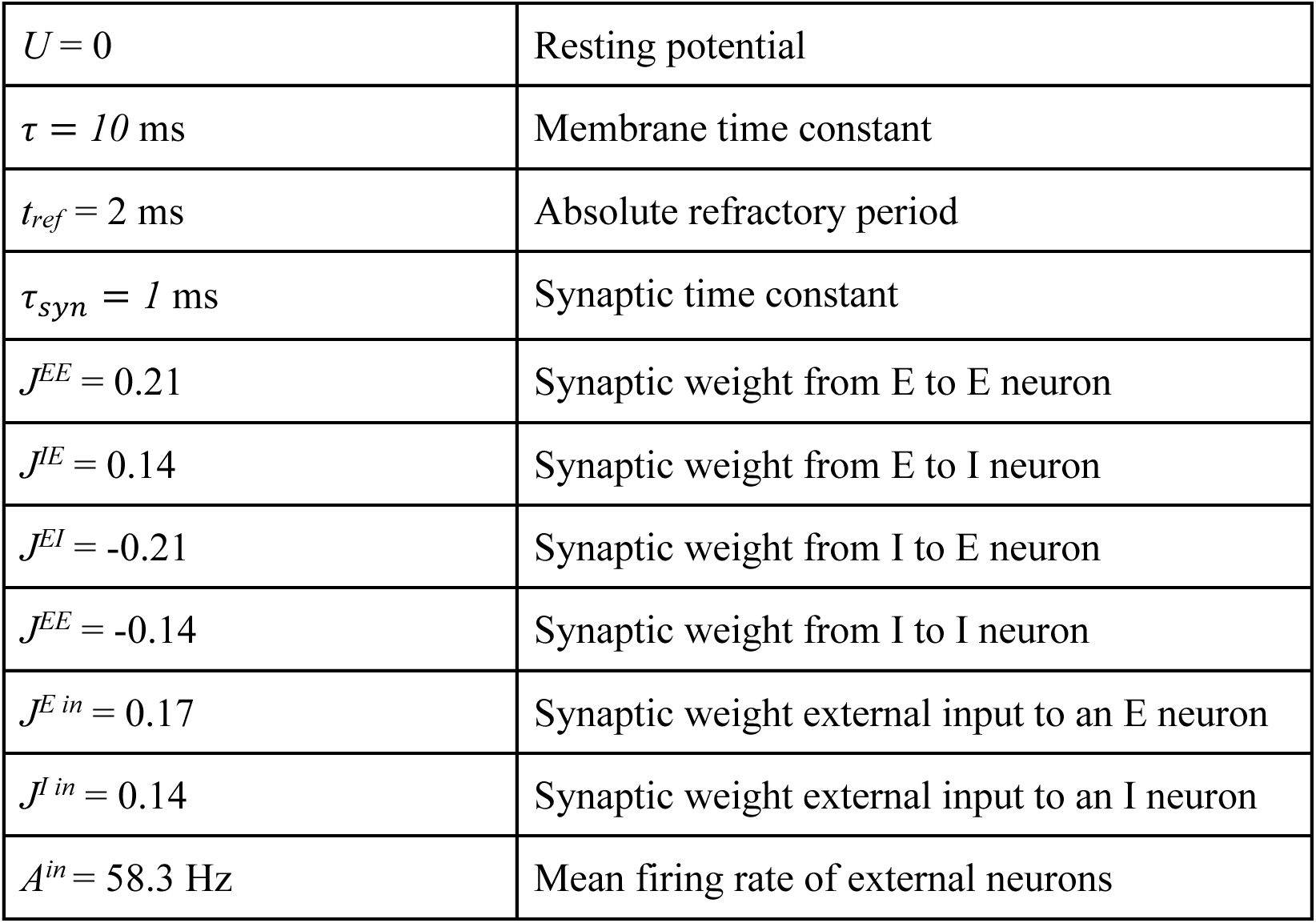

The network is divided into excitatory and inhibitory populations of size *N^E^* and *N^I^* respectively, with excitatory neurons constituting 80% of the population. The connectivity of the network is sparse and generated by sampling each connection as an independent Bernoulli variable with probability 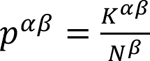 where α ∈ {E, I} and β ∈ {E, I, Ext}. Nonzero synaptic weights between two populations all have the same value, i.e. if J^αβ^_ij_ is nonzero then 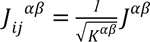 where α, β ∈ {E, I} (see **Table 1**). The average number of incoming per neuron was independent of the neuron type, i.e. *K*^EE^ = *K*^EI^ = *K*^IE^ = *K*^II^ = *100*. To vary the sparsity of the network connectivity, we varied the number of neurons in the network while keeping the number of incoming and outgoing connections constant. In this way, we generated networks with connection probabilities *p*^EE^ ∈ {*2*.*5%*, *5%*, *10%*, *12.5%*, *25%*}. Notice that connection probabilities from inhibitory neurons were 4 times larger.

We simulated 3 synthetic ground truth networks for *T* = 14,400 seconds of simulated time for each sparsity level. We then randomly sampled 3 sets of 100 inhibitory and 400 excitatory neurons to generate synthetic ground truth datasets of connectivity and activity for each network realizations.

### Assessing Dyad’s precision using putative neuron type

To estimate Dyad’s precision in identifying pairs connected by an excitatory synapse in our own data, we took advantage of the fact that narrow-spike, high-firing rate units are overwhelmingly inhibitory^27,66–70^. We therefore obtained a conservative estimate of Dyad’s precision by assuming that any detected excitatory connection outgoing from one such neuron was a false positive.

For this analysis, we considered only units that had a narrow waveform (trough-to-peak distance < 0.5ms) and firing rate larger than 5Hz. We then counted how many outgoing excitatory connections from these units were detected by Dyad (2), resulting in a false-positive rate (*FPR*) of 1.3 x 10^-5^. To obtain an estimate of precision, we then used the expression:

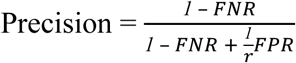

where *FNR* and *FPR* are the false-negative and false-positive rate, respectively, and *r* is the class imbalance i.e. the ratio of positive over negative pairs in the dataset. Thus, to obtain the precision we had to assume a value for the *FNR* (note that *r* can be derived from our measurements given *FPR* and *FNR*). However, the precision estimate depended very weakly on *FNR* and was approximately 0.98 for most values of the *FNR*.

We used an analogous approach to estimate the precision of Dyad in detecting neuron pairs connected by an inhibitory synapse. Because we could not use spike waveform features or firing rate to identify a population of neurons that are overwhelmingly excitatory, we considered as false positive any pair having an inhibitory connection outgoing from an E neuron for which Dyad identified an excitatory connection as well. We obtained a *FPR* of 9.0 x 10^-6^ (13 false positives out of 1,448,637 potential pairs). Using the above expression to relate *FPR* to the precision, we obtained a precision > 0.99 when considering only I neurons as pre-synaptic, for a broad range of *FNR*. We note that if we were to consider all possible neurons as pre-synaptic the dataset would be much more imbalanced (smaller *r*) and the precision would be lower (0.93).

### Assessing Dyad’s precision using CCG peaks on anti-causal side

We also estimated Dyad’s precision by measuring how many times our pipeline would detect synapses on the anti-causal side of the CCG. Indeed, peaks induced by confounding factors are not necessarily on the causal side (in contrast to those caused by synaptic connections). We used Dyad (without retraining) to rank CCGs based on an anti-causal window that extends from −15 to 5 milliseconds. Because this window is shifted by 5 milliseconds compared to the one used to train Dyad, it will tend to look for peaks between −4.5 and −1.5 millisecond lags. Applying the same criteria for synaptic connectivity as described above, we found only 12 false positives, yielding an estimated *FPR* of 8.1 x 10^-6^. Using the same method described above, we obtain an estimate of the precision above 0.98.

We used the same approach to estimate the precision of the inhibitory version of Dyad. In this case, we detected 7 false positives out of 1,490,149 (N = 2 mice) potential interactions, resulting in an estimated *FPR* of 4.7 x 10^-6^. Using the method described above to relate *FPR* to the precision, we obtained a precision > 0.99 when considering only I neurons as pre-synaptic, for a broad range of *FNR*. If instead we were to consider all possible neurons as pre-synaptic the dataset would be more imbalanced, resulting in a lower precision (0.97).

### Assessing the Effect of Disynaptic Chains

To estimate the extent to which the excitatory CCG peaks detected by Dyad might reflect disynaptic chains instead of monosynaptic connections, we took advantage of the units for which Dyad had identified at least one incoming and one outgoing connection. In these cases, we had that neuron *i* connects to neuron *j* which connects to neuron *k*, resulting in a disynaptic chain from *i* to *k*. We then asked whether Dyad detected a connection from *i* to *k* more than expected based on a null model. To take into account the fact that some neurons make more connections than others, we constructed a null model that preserved the number of incoming and outgoing connections of each neuron but shuffled the identity of pre and postsynaptic partners.

### Identification of E and I Neurons

Inhibitory neurons are characterized by thin spike waveforms and high baseline firing rates^27,66–70^. To identify putative excitatory and inhibitory neurons, we trained a support vector machine classifier with polynomial kernel (degree = 5) on two features of individual units: the firing rate and the waveform trough-to-peak distance. To train the classifier, excitatory (inhibitory) labels were assigned based on whether the unit was the presynaptic neuron in an excitatory (inhibitory) connected pair identified by Dyad. The trained SVM identifies a nonlinear boundary separating E from I neurons in the trough-to-peak – firing rate plane (**Extended Data** Fig. 1h). Such boundary reflects the fact that inhibitory neurons tend to have narrower waveform and higher firing rate, both of which are known physiological properties of the majority of inhibitory neurons in the piriform cortex^67–70^.

We found 189 I neurons out of 5,478 total single units (3.5% of the population), consistent with the low fraction of inhibitory neurons in the piriform^69,71,72^. We note that the population we define as E neurons contains a small minority of inhibitory cells with broad spike waveforms, which likely correspond to somatostatin-positive neurons^70^. We then identified superficial Layer 1 feedforward inhibitory neurons in the I neuron population by the estimated spatial location. These cells receive input from the olfactory bulb but negligible recurrent input from cortex^73^ and were excluded from further analysis. The remaining I neurons, which were included in our analyses, correspond predominantly to feed-back interneurons and receive little to no direct input from the olfactory bulb^73^.

The E-to-E connection probability was 0.032% (Fig. 1b), lower than the expected 0.1% and consistent with our network simulations in which Dyad recovers only a subset of the connected pairs of the network. The E-to-I and I-to-E connection probabilities were higher: 1.1%, and 0.81% respectively (Fig. 1b). As we only detected 8 I-to-I connections (0.13%) we did not consider these further. The relatively elevated connection probability of E-to-I compared to E-to-E is consistent with a previous report of extensive excitatory convergence onto local inhibitory neurons in the piriform^73^. We note that this higher E to I connectivity may also reflect a detection bias favoring the relatively high firing rates and input resistances of inhibitory neurons.

### Computational Model

We developed a simplified model to study the effect of inhibitory rewiring on the odor responses of excitatory neurons in the piriform cortex. In the model, the responses of N_I_ = 2,000 inhibitory neurons to odorant stimuli are sampled from a log-normal distribution independently for each neuron but with correlations among different odors, i.e.

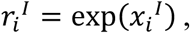

Where *r*_i_^I^ is a vector of length *P*, the number of odors, containing the responses of inhibitory neuron *i* to the odorant stimuli. *x*_i_^I^ is a vector of the same length sampled from a multivariate normal distribution with mean zero and covariance matrix

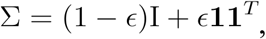

where I is the *P*-dimensional identity matrix and **1** is a length-P vector of all ones. The parameter ε controls the strength of correlations among different odorant stimuli. For all the analyses, we set ε = *0*.*6* and P = 8. Among the eight simulated stimuli, the first four were assumed to be the familiar and the second four the novel ones.

We considered two models for generating the connectivity matrix *W* between inhibitory and excitatory neurons. In the first model we sampled the probability of forming outgoing connections *p*_i_^*out*^ from a log-normal distribution with mean 0.05 and variance 0.0095, independently for each inhibitory neuron. Synaptic weights from inhibitory neuron *i* to excitatory neurons are either equal to −1 with probability *p*_i_^*out*^or 0 otherwise. We refer to this model as naïve, because there is no relationship between the probability of inhibitory neurons to form outgoing connections and their odor responses.

The second model was obtained from the first by assigning the previously sampled output probabilities based on the sampled odor responses of the inhibitory population: inhibitory neurons were ranked based on their selectivity index (S.I.) for the set of familiar odors and were assigned the output probabilities in decreasing order based on such ranking. We note that because of the stochasticity of the process that generates connections from output probabilities, it is not certain that inhibitory neurons with the largest S.I. will form the most output connections.

In addition to input from inhibitory neurons, excitatory neurons received another source of input representing other potential sources of inputs such as the olfactory bulb, recurrent excitation from the piriform, and input from other brain regions. Such input was sampled from the same distribution as the inhibitory neuron responses, i.e. log-normal with correlations among odorant stimuli, but in this case independently for each excitatory neuron. The strength of such input was set to 0.3 times the strength of the inhibitory input. To account for trial-by-trial variability, we added 25 independent Gaussian noise realizations to both the inhibitory responses and the external input, independently for each neuron, resulting in 25 trials. The strength of such noise was set to 20% of the signal that it was applied to.

To summarize, the total input excitatory neuron *i* in response to odor μ in a given trials was

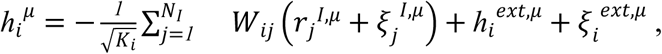

where *h_i_^ext^* is the external input, ॐ^I^ and ॐ^ext^ are the inhibitory and external response noise. The factor 1/√K_i_, where *K_I_* is the in-degree of excitatory neuron *i*, was added to make the variance of the total inhibitory input of order one^74^. Finally, to mimic a recurrent balance mechanism in our simplified model, we assumed that for each odor the total input current to excitatory neurons was on average zero^75^. The odor responses of excitatory neurons were obtained by passing the balanced input *h*^u^_i_ through a ReLU nonlinearity, with threshold θ = *2*.*5*. All our results were obtained using the responses of 500 model excitatory neurons and averaged across 100 realizations of the model.

### Data Analysis

We defined the response of a neuron to an odor in a trial as its number of spikes in a two-second window after the time of valve opening. We found that our results were robust when the length of this window was changed between 1 and 4 seconds (**Extended Data** Fig. 7b, not shown for other results). Most of our analyses were based on trial-averaged responses. However, for measuring cross-validated signals correlations, coefficient of variation across trials, and classifications performance we used odor responses in individual trials.

### Unit Location

As previously described^62^, briefly: We computed a spatial average across electrode site locations, estimating each single unit’s position (x,y) as:

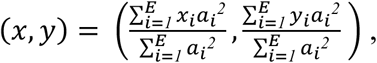

where *E* is the number of electrode sites, *x_i_* and *y_i_* are the lateral and vertical position of electrode site *i*, and *a_i_* is the peak-to-peak amplitude of the mean spike waveform recorded at the electrode site *i*. The choice to square the peak-to-peak waveform amplitude was to mitigate the effect of very low-amplitude waveforms registering on all of the probe’s electrode sites.

### Signal Correlations

Signal correlations between two neurons were measured using the Pearson’s correlation coefficient between the odor responses. To mitigate the problem of spurious correlations arising from the small number of odors considered, we used a cross-validated approach. The correlation coefficient was measured based on responses averaged across a different subset of trials for the two neurons. This operation was repeated for five cross-validation folds, and the resulting correlation coefficients were averaged to get a cross-validated quantity. When computing signal correlations across only familiar or novel odors the number of cross-validation folds was increased to ten to obtain a distribution of signal correlations with the same support as when computing signal correlations across all 8 odors. All our results are robust to varying the number of cross-validation folds and hold when using standard, non cross-validated correlation coefficients as well.

### Lifetime and Population Sparseness

We defined the population sparseness of an odor α measured across a population of *N* neurons as

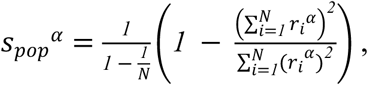

Where *r*_i_^α^ is the response of neuron *i* to odor stimulus α.

Similarly, the lifetime sparseness of a neuron measured across a set of *P* odors was

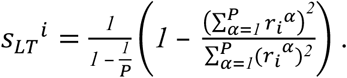

### Spatially-structured null model of like-to-like connectivity

We separated all pairs of simultaneously recorded units in our dataset in 50 groups based on the estimated unit distance, and measured the connection probability for each group. The distance bins were set in such a way to have an equal number of pairs in each bin. We then used these quantities to generate a null model in which the observed spatial organization of the piriform was preserved, i.e. nearby pairs were more likely to be connected and slightly more correlated than more distant ones, but connection probability was independent of response properties. For each realization of this null model, we sampled each connection independently based on the pair distance; we sampled the pair signal correlation also based on the distance by sampling with replacement from the correlations of the pairs in the same distance group. For each null-model realization we measured the relationship between connection probability and signal correlation across the population, and computed statistics across 100 null model realizations.

### Single neuron selectivity In Extended Data Fig 2x

we quantified single neuron odor selectivities using decoding performance on a trial-by-trial basis. For each neuron and each odor, we trained a linear SVM to distinguish responses to the odor from the other stimuli using a one-vs-one scheme and leave-one-out cross-validation across trials. This approach yielded eight numbers for each neuron, corresponding to the average classification performance of each odor stimulus against the others. The maximum selectivity quantifies the selectivity of a neuron to the odor it was most selective to and was then obtained for each neuron by taking the maximum of the classification performance across odors.

### Predicting connectivity from changes in single-neuron response properties

To investigate whether single-neuron properties might affect connectivity in an experience-dependent manner, we asked whether there was a significant relationship between the number of incoming or outgoing connections and the indices

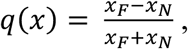

Where *x* is a scalar function of single-neuron response properties that can be computed for either novel (*x_N_*) or familiar (*x_F_*) responses.

We tested three choices for *x:* the odor response, averaged across odors in a set; the coefficient of variation of the response across trials, and the standard deviation of the response across odors in a family. The significance of a relationship between *q* and in- or out-degrees was assessed using Pearson’s correlation coefficients, although our results also hold when considering Spearman’s correlations instead. In total, we tested three connectivity features (in-degree of E neurons, out-degree of E neurons, in-degree of I neurons) against three possible predictors, resulting in nine comparisons. To correct for multiple comparisons artifacts, we applied the Bonferroni correction.

### Dimensionality of Odor Responses

The dimensionality of excitatory neuron odor responses was measured by the participation ratio of the eigenvalues λ of the neuron-by-neuron covariance matrix of odor responses^47^, i.e.

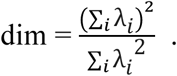

### Odor Response Discriminability

We assessed the discriminability of different odorant stimuli based on the odor responses using a linear classifier on single-trial responses. To avoid that the classifier accuracy approached 1, making comparisons between novel and familiar odorant stimuli more difficult, we measured the classifier accuracy as a function of the number of neurons considered, ranging from 15 to 300, in intervals of 15. For each of these values, we considered 200 random subsets of neurons. For each subset, trials were randomly separated into training and testing using two-fold stratified cross-validation. Multi-class accuracy was assessed using a one-versus-rest approach and the mean and variance of the classifier performance was measured across odors, cross-validation folds, and neuron subsets.

We used a Hebbian classifier, which mimics a downstream neuron which receives input from the excitatory population and can learn to discriminate odors based on a biologically-plausible Hebbian rule. In this classifier, readout weights are set by

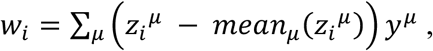

Where *y*^μ^ is the label of odor μ and *Z*_i_^μ^ is the response of the excitatory neuron *i* to odor μ averaged and z-scored across all trials in the training set. Test accuracy was then assessed by computing the predicted labels as

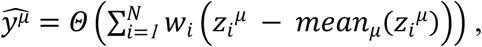

where Θ is the Heaviside step function and *Z*_i_^μ^ was computed using trials in the test set. Similar results were obtained using a linear support vector machine classifier, although the difference between novel and familiar stimuli was reduced.

### Estimation of the Effect of Experience on Inhibitory Currents

We used both measured inhibitory responses and connectivity to estimate the effect of the observed experience-dependent organization on total inhibitory currents onto excitatory neurons. First, because the number of inhibitory neurons per dataset was relatively small, we concatenated all the inhibitory neurons, forming an inhibitory pseudopopulation. Each inhibitory neuron in the data was then assigned a probability of forming outputs to excitatory neurons based on the measured number of detected outputs 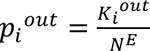, where *N^E^* is the number of excitatory neurons in the dataset in which inhibitory neuron *i* was recorded.

These probabilities were used to sample the connections *W*_i_ between the population of inhibitory neurons and an example excitatory neuron as independent Bernoulli variables. Because excitatory neurons in the piriform can receive inputs from a much larger pool of inhibitory neurons than the recorded ones, using the measured probability might result in undesired effects due to small-sample-size. To prevent this, we scaled all connection probabilities by a factor 10. The resulting connectivity was then used to estimate the inhibitory input current *ℎ*^I^ onto the excitatory neuron, assuming that currents from different neurons sum linearly and that all synaptic efficacies are equal:

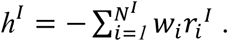

